# Cytokine-bearing Bacterial Outer Membrane Vesicles with Empowered Efficacy in Intratumoral Immunotherapy

**DOI:** 10.64898/2026.04.02.716109

**Authors:** Riccardo Corbellari, Michele Tomasi, Mattia Benedet, Assunta Gagliardi, Rubens Begaj, Ilaria Zanella, Silvia Tamburini, Elena Caproni, Enxhi Shaba, Gabriele Di Lascio, Valeria Facchini, Chiara Baraldi, Gaia Gambini, Alvise Berti, Andrea Lunardi, Luca Bini, Guido Grandi, Alberto Grandi

## Abstract

Bacterial Outer Membrane Vesicles (OMVs), spherical bilayered nanoparticles naturally released by all Gram-negative bacteria, are gaining increasing interest not only in the design of prophylactic vaccines but also in cancer immunotherapy. In particular, thanks to their potent built-in adjuvanticity and to their intrinsic capacity to directly kill tumor cells, OMVs have been successfully tested in intratumoral *in situ* vaccination (ISV), a strategy in which immunostimulatory formulations are injected directly into tumors to convert the tumor microenvironment (TME) into an immune-reactive state. Previous studies have shown that OMVs induce robust inflammation and a Th1-skewed immune response, resulting in complete tumor remission in a substantial fraction of mice bearing syngeneic tumors. Here, we show that OMVs from our *Escherichia coli* Δ60 strain can be efficiently engineered with multiple cytokines and chemokines. Moreover, CCL3, Flt3L, TNFα, and IL-2 not only accumulated on the OMV surface but also retained their *in vitro* biological activity. Furthermore, OMVs displaying these cytokines exhibited potent antitumor activity, and in particular the intratumoral injection of the combined TNFα- and IL-2-engineered OMVs eradicated tumors in over 95% of mice across several syngeneic models. Immunostaining and flow cytometry analyses revealed that injection of engineered OMVs markedly remodeled the TME, promoting the recruitment of inflammatory myeloid cells and γδ T cells, the persistence of local CD8⁺ and CD4⁺ αβ T cells, and the reduction of regulatory T cells. Overall, these results highlight cytokine-bearing OMVs as a versatile and highly effective platform for intratumoral immunotherapy.

## INTRODUCTION

*In situ* vaccination (ISV) is a cancer immunotherapy strategy which envisages the direct injection of immunostimulatory reagents into the tumor mass. The rationale stems from the evidence that the immune system fails to recognize and kill cancer cells in full blown tumors because of the strong immunosuppressive tumor microenvironment. To overcome this hurdle, tumors are inoculated with different medicinal recipes, all aiming at reverting the microenvironment into an immune-reactive one (*1*).

The first reported practice of ISV is likely found in an ancient Egyptian scroll describing how the renowned physician Imhotep (ca. 2600 BC) recommended treating tumors (swellings) with a poultice followed by incision (*2*). Such a regimen would inevitably have led to infection and, in some cases, tumor elimination. At the end of the nineteenth century, Dr. William Coley, observing several patients who fully recovered from cancer when an infection occurred at the surgical sites, developed a variety of strategies for treating cancers with live and dead bacteria or with bacterial extracts, later named “Coley’s Toxin” (*2,3*). He treated 896 patients, achieving five-year survival rates of 34% to 73% for inoperable carcinomas and of 13% to 79% for inoperable sarcomas. The Coley’s toxin is still used in Germany and, importantly, paved the way to the development of the tuberculosis *Bacillus Calmette-Guerin* (BCG) vaccine, approved in 1990 by the Food and Drug Administration (FDA) for the treatment of intermediate- and high-risk non-muscle invasive bladder cancer (NMIBC) (*4*). Nowadays, in addition to BCG, three ISV therapies have received official approval in different countries. In 2015, the oncolytic virus talimogene laherparepve (T-VEC) was approved both in the USA and EU for advanced melanoma patients (*5*). In 2019, Hensify® (also known as NBTXR3) was approved in Europe for the treatment of soft tissue sarcoma in combination with radiotherapy (*6*). Finally, in 2021, Japan approved DELYTACT® (teserpaturev/G47Δ), a genetically engineered oncolytic herpes simplex virus type 1 (HSV-1), for the treatment of malignant glioma (*7, 8*).

Interest in ISV is steadily increasing, as evidenced by the growing number of published pre-clinical and clinical studies (*9*) and by the 133 clinical trials that, as of April 1^st^, 2026, are listed on ClinicalTrials.gov under the search term ‘in situ immunotherapy’.

Although all ISV formulations share the common goal to turn the tumor microenvironment into an immune reactive one, they differ in how they attempt to do so, each formulation privileging specific mechanisms of anti-tumor immune response (*9*). For instance, the TLR3, TLR4, TLR7/8, TLR9 and STING agonists, used in 25% of the current clinical studies, predominantly act as “Th1 adjuvants”, recruiting and activating NKs, neutrophils, M1 macrophages, T cells and DCs specialized in cross-presentation. The oncolytic viruses (up to 40% of all reported clinical trials) cause immunogenic cell death (ICD) of cancer cells thus promoting the spreading of tumor antigens. Finally, tumor-antigen specific mAbs (10% of the current studies) kill cancer cells via ADCC/CDC/toxin delivery while checkpoint inhibitor antibodies activate cytotoxic CD8^+^T cells and inhibit CD4^+^ T_REG_. Ideally, to be highly effective, ISV should trigger, in a concerted manner, different mechanisms of anti-tumor immune responses.

In line with this concept, we have recently explored the use of bacterial Outer Membrane Vesicles (OMVs) for ISV. We hypothesized that OMVs, spherical, bilayered nanoparticles naturally released by Gram-negative bacteria through outer-membrane budding (*10*), could be appealing for ISV because they can activate two major immune mechanisms. First, OMVs possess intrinsic adjuvanticity due to multiple microbe-associated molecular patterns (MAMPs) present in the outer membrane and periplasm (e.g., LPS, lipoproteins, peptidoglycan), which trigger strong inflammation and a Th1-skewed response (*11*). Second, OMVs from several bacterial strains, including *E. coli*, can directly kill tumor cells by a mechanism known as pyroptosis (*12, 13*), thereby enhancing antigen release within the tumor microenvironment. Consistent with these properties, we showed that intratumoral OMV injection elicited robust anti-tumor responses in syngeneic mouse models, achieving complete tumor eradication in 40–60% of animals depending on the model (*14*). Encouraged by these results, we asked the question as to whether the therapeutic efficacy of OMVs could be further improved. We reasoned that if we could engineer the OMVs with selected cytokines/chemokines we would combine three mechanisms of anti-tumor immune responses in a single therapeutic agent.

Cytokines are potent regulators of immune activation (*15*), and several classes—including interleukins, interferons, TNFs, chemokines, and growth factors—already play important roles in cancer immunotherapy, with some approved for clinical use (*16–19*). Cytokines are also being evaluated in ISV with promising results (*1, 9*). Among those tested in ISV, Flt3L, CCL3, TNFα, and IL-2 have shown notable therapeutic activity. Flt3L recruits BATF3^+^ dendritic cells, which are crucial for generating tumor-specific cytotoxic T cells, and is under evaluation in multiple clinical trials (*20, 21*). CCL3 recruits diverse immune cells, correlates with improved prognosis (*22, 23*), and has shown preclinical efficacy in ISV (*24*). TNFα, produced mainly by myeloid cells but also by activated T cells and stromal cells, enhances acute anti-tumor responses (*18*). IL-2, the first cytokine ever used in cancer immunotherapy, promotes T cell activation and T cell and NK cell expansion and is approved for metastatic RCC and melanoma (*18*). Importantly, the combination of TNFα and IL-2 in the targeted biopharmaceutical Nidlegy® (Daromun) has demonstrated potent efficacy in Phase II trials for stage IIIB/C melanoma, BCC, and SCC (*25, 26*), and is currently under evaluation in Phase III studies (NCT02938299, NCT03567889).

In this work, we first show that OMVs from our *E. coli Δ60* strain (*27*) can be efficiently decorated with diverse cytokines and chemokines. Next, we focus on CCL3, Flt3L, TNFα, and IL-2, demonstrating that these cytokines are robustly expressed and properly displayed on both bacterial and OMV surfaces, while retaining their biological activity. Finally, we show that OMVs engineered with these cytokines exhibit enhanced anti-tumor efficacy *in situ*, with the TNFα-OMVs + IL-2-OMVs combination (Combo) completely eradicating tumors across multiple syngeneic mouse models.

## RESULTS

### OMVs from E. coliΔ60 can be efficiently engineered with a variety of cytokines

Our strategy to express cytokines in the OMVs envisages their fusion to the C-terminus of the *Staphylococcus aureus* FhuD2 lipoprotein. We previously showed that FhuD2 can be efficiently expressed on the surface of *E. coli* using the lipoprotein transport machinery. Moreover, the protein accumulates in the OMVs, where it represents more than 20% of the total OMV proteins (w/w) (*28*). We also previously showed that FhuD2 can be conveniently exploited as a carrier to transport a variety of small proteins, protein domains and polypeptides to the vesicular compartment (*29*). Thanks to this background, we created recombinant expression plasmids in which the synthetic genes encoding a set of cytokines were fused in frame to the 3’ end of the FhuD2 gene (Figure 1A). The plasmids were used to transform *E. coli*Δ60 strain (*27*), generating recombinant clones each expressing a specific FhuD2-cytokine fusion. The recombinant strains were grown in culture and the OMVs were purified from the culture supernatants and analyzed by SDS-PAGE. As shown in Figure 1B, all fusion proteins efficiently accumulated in the vesicular compartment, representing from 10 to 15% of total OMV proteins (w/w) as judged by densitometric scanning of the Coomassie Blue-stained gel. We also investigated the total proteins present in TNFα-OMVs and IL-2-OMVs by 2D electrophoresis. For this analysis, each OMV preparation was run on three 2D gels to carry out proper statistical analysis. As shown in Figure S1A, the 2D maps of the two OMV preparations were essentially superimposable with the exception for six protein species that were specific for each preparation (see arrows in Figure S1A). The similarity between the two OMV preparations is clearly highlighted in Figure S1B, which shows the Principal Component Analysis (PCA) analysis of the six 2D maps. The PC1 values are highly similar and the statistical analysis clusters the three 2D maps of each preparation in two distinct groups. To characterize the six unique proteins present in each OMV preparation, we carried out a 2D Western Blot analysis using IL-2- and TNFα-specific antibodies. As shown in Figure S1C, the anti-IL-2 antibodies stained the six protein spots exclusively present in the IL-2-OMV 2D map. Likewise, the anti-TNFα antibodies recognized the six protein spots specific for the TNFα-OMVs. Overall, these data demonstrate that the recombinant OMVs are highly homogeneous in terms of protein content, the only major difference being the recombinant protein itself.

**Figure 1.**
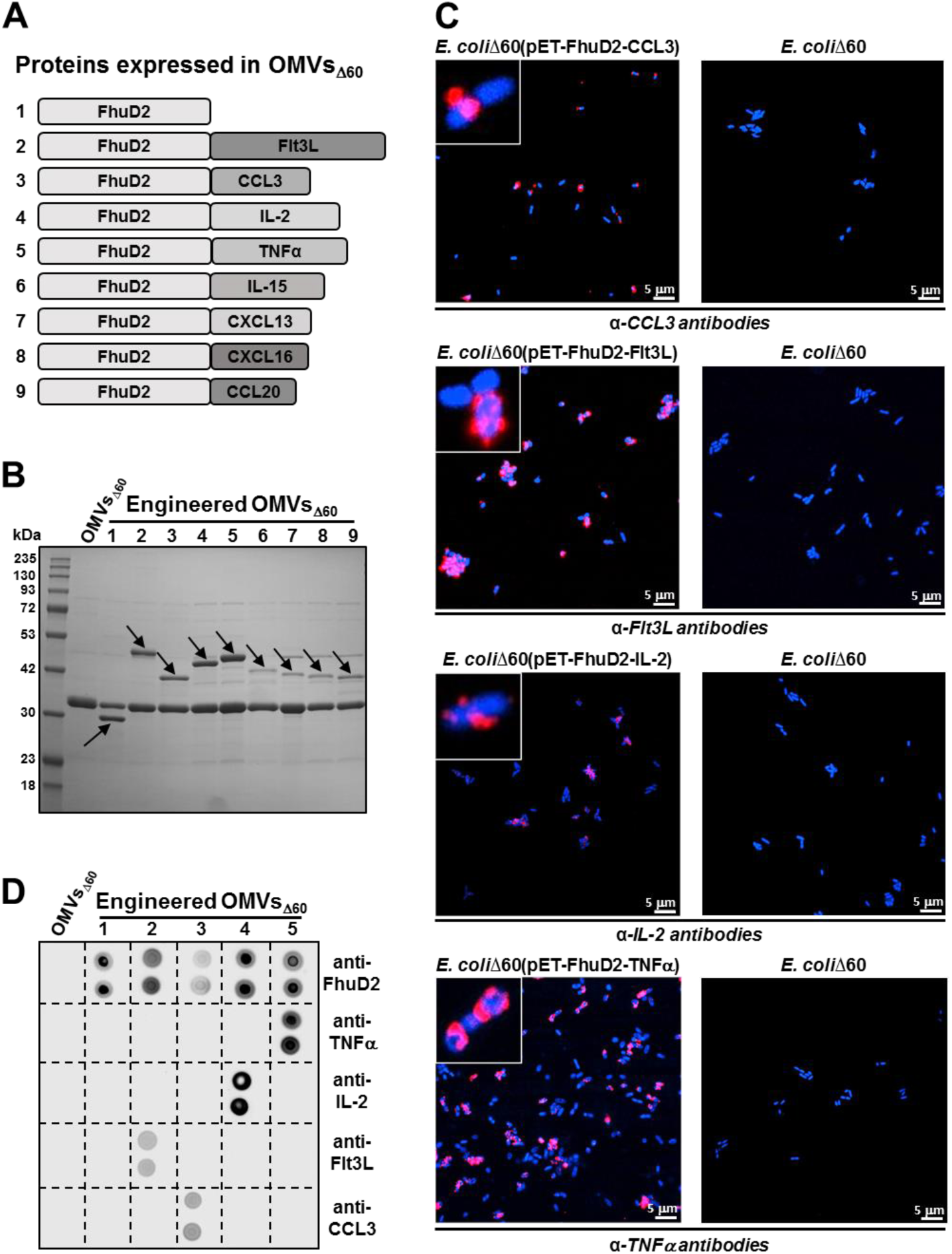
Expression and localization of cytokine fusions in recombinant E. coliΔ60 strains and in engineered OMVs_Δ60_. **(A)** Schematic representation of cytokine fusions – Genes encoding the different cytokines were fused to the 3’ end of the *Staphylococcus aureus* FhuD2 gene. The fusions were inserted into the pET plasmids and the plasmids were used to transform the OMV-overproducing strain *E. coli*Δ60. (**B**) *SDS-PAGE analysis of OMVs purified from engineered E. coliΔ60 strains* – OMVs (15 μg) purified from the supernatants of each engineered strain and were analyzed by SDS-PAGE. (**C**) *Confocal microscopy analysis of recombinant E. coliΔ60 strains expressing CCL3, Flt3L, TNFα and IL-2 fusions*. *E. coli*Δ60 cells expressing the cytokine of interest and *E. coli*Δ60 cells as a negative control were first incubated with primary anti-cytokine mouse antibodies and subsequently with Alexa Fluor-conjugated anti-mouse antibodies. Samples were mounted on microscopy slides and examined by confocal microscopy. (**D**) *Dot blot analysis of engineered OMVs* - Flt3L-OMVs_Δ60_, CCL3-OMVs_Δ60_, IL-2-OMVs_Δ60_, TNFα-OMVs_Δ60_ and “Empty”-OMVs_Δ60_ were spotted in duplicate onto a nitrocellulose membrane (2 μg) and membranes were incubated with the cytokine-specific primary antibody. As a control, the primary antibody against the FhuD2 carrier protein was also tested. The filters were subsequently incubated with specific HRP-conjugated secondary antibodies and antibody binding was detected with the ImageQuant LAS4000 using the SuperSignal West Pico chemiluminescent substrate.

### OMV-associated Flt3L, CCL3, TNFα and IL-2 are surface expressed and preserve their biological function

Having demonstrated that cytokines compartmentalize in the vesicles as FhuD2 fusions, we next focused our attention on the four recombinant strains *E. coli*Δ60(pET-FhuD2-Flt3L), *E. coli*Δ60(pET-FhuD2-CCL3), *E. coli*Δ60(pET-FhuD2-TNFα) and *E. coli*Δ60(pET-FhuD2-IL-2).

To assess whether these cytokines protruded out of the membrane, we analyzed the recombinant strains by confocal microscopy using cytokine-specific antibodies. As shown in Figure 1C, non-permeabilized bacteria were efficiently stained by the antibodies recognizing the cytokine expressed by each strain.

Next, to reveal the presence of the fusion proteins on the surface of the engineered OMVs, we spotted aliquots of the four engineered OMVs (Flt3L-OMVs_Δ60_, CCL3-OMVs_Δ60_, TNFα-OMVs_Δ60_ and IL-2-OMVs_Δ60_) on a nitrocellulose membrane and we analyzed the capacity of the corresponding anti-cytokine antibodies to bind to the immobilized vesicles. As shown in Figure 1D, the Dot Blot analysis indicates that the cytokine-specific antibodies recognized the immobilized vesicles in a highly specific manner. As a control, we also used antibodies recognizing a FhuD2 epitope localized at the N-terminus of the protein. Such antibody reacted with all cytokine-expressing OMVs, indicating that the cytokines well protruded out of the membrane vesicles, being the FhuD2 carrier protein accessible to the binding of its specific antibodies.

We next asked the question whether, when immobilized to the OMV surface, the four cytokines were still biologically active. To address this question, *in vitro* cell assays were carried out. In the case of CCL3, the PBMC migration assay was used. PBMCs isolated from mouse blood by Ficoll gradient were seeded in the upper chamber of a cell migration assay device set with a 5.0 µm polycarbonate membrane. The addition of OMVs to the lower chamber promoted the migration of a substantial fraction of PBMCs towards the OMVs (Figure 2A). The biological activity of the OMV-associated IL-2 was evaluated using the cytotoxic T lymphocytes proliferation assay. CTLL-2 cells (2.5 × 10⁴ in RPMI culture medium) were exposed to serial dilutions of IL-2-OMVs and cell viability was analyzed after 72-hour incubation at 37°C. As shown in Figure 2B, a dose-dependent cell proliferation was observed. As far as the activity of the OMV-associated TNFα is concerned, MC38 carcinoma cells (2 × 10⁴ cells per well) were incubated with different amounts of TNFα-OMVs and cell survival was assessed after 24 hours. As shown in Figure 2C, the presence of TNFα in the vesicles increased the natural capacity of OMVs to kill cancer cells in a statistically significant manner. Finally, regarding Flt3L, we could not confirm its *in vitro* activity when associated with the OMVs since OCI-AML5, the cells commonly used to measure Flt3L proliferative property, are particularly sensitive to the killing by OMVs. In the absence of data demonstrating the functional activity of OMV-immobilized Flt3L, before moving to *in vivo* experiments (see below) we wanted to know whether Flt3L was still functional when fused to the C-terminus of a carrier protein. To this aim, we expressed the MBP-Flt3L fusion, which was subsequently purified by affinity chromatography. Next, the activity of the purified protein was tested *in vitro* and, as shown in Figure 2D, MBP-Flt3L promoted OCI-AML5 cell proliferation in a dose-dependent manner.

**Figure 2.**
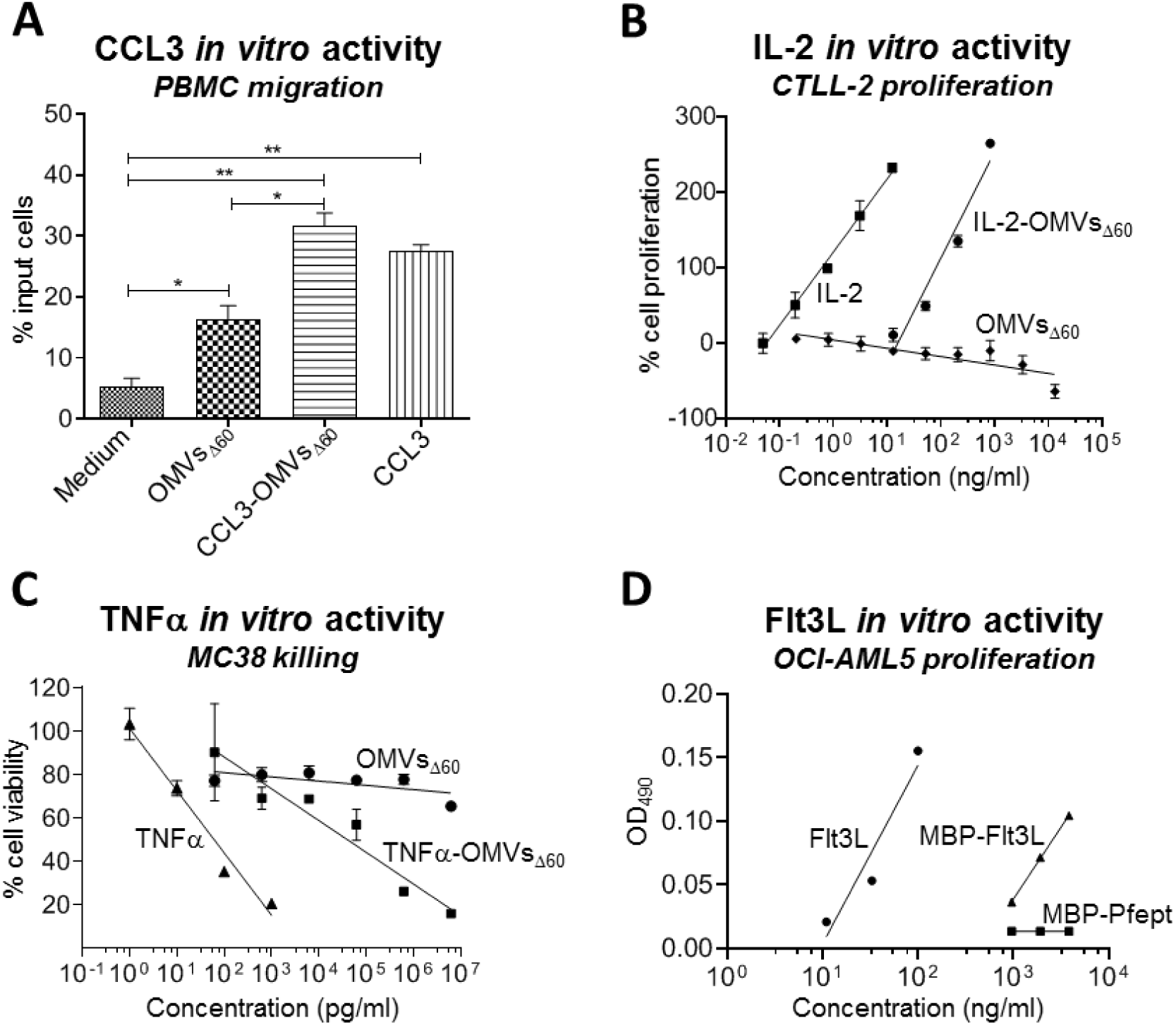
In vitro activities of OMV-associated cytokines. (**A**) *CCL3-OMVs_Δ60_ in vitro activity* - Mouse PBMCs (1.5 × 10⁵) were seeded in the upper chambers of a Corning® Transwell® apparatus equipped with a 5.0 µm polycarbonate membrane. The lower chambers contained RPMI alone or supplemented with OMVs_Δ60_ (1 µg), mCCL3-OMVs_Δ60_ (1 µg), or recombinant mCCL3 (30 nM). Cells migrating to the lower chamber were counted using a Neubauer chamber. Statistical analysis was performed using an unpaired *t*-test. (**B**) *IL-2 in vitro activity **-*** The biological activity of the OMV-associated cytokine IL-2 was determined by its ability to stimulate the proliferation of murine cytotoxic T lymphocytes (CTLL-2). CTLL-2 cells were seeded in a 96-well plate and then serial dilutions of OMVs_Δ60_, IL-2-OMVs_Δ60_ or recombinant IL-2 were added to the plate. Cell proliferation was determined after incubation at 37°C for 72 hours. (**C**) *TNFα in vitro activity -* MC38 cells (2 × 10⁴ cells per well) were incubated 24 hours with serial dilutions of either OMVs_Δ60_, TNFα-OMVs_Δ60_ or recombinant TNFα. Cytotoxicity was determined after 3 hours and expressed as the percentage of viable cells with respect to control (cells treated with actinomycin D only). (**D**) *In vitro activity of MBP-Flt3L fusion-* The biological activity of MBP-Flt3L fusion was determined by measuring its ability to stimulate the proliferation of OCI AML-5 cells. The cells, grown in Alfa-MEM medium supplemented with 20% FBS + 10 ng/ml GM-CSF, were dispensed into a 96-well plate (1.2 x 10^5^ cells/well), and stimulated for 48 hours at 37°C with either recombinant Flt3L, or MBP-Flt3L, or MBP-Pfpep (negative control).

To investigate the activity of the OMV-associated cytokines in more quantitative terms, we also ran each functional assay using the corresponding purified recombinant proteins and we compared the activity of the immobilized cytokines (attached to the OMVs) with the activity of their soluble forms (Figure 2A-D). To make the quantitative comparison, using densitometric analysis of OMV protein bands separated by SDS-PAGE, we estimated that the amount of each cytokine in the OMVs was approximately 25 ng/μg of OMVs (each fusion protein corresponds to approximately 10% of the total OMV proteins (w/w) and cytokines represent approximately one fourth of the fusions (10 kDa out of 40 kDa)). Regarding CCL3, 1μg of CCL3-OMVs and recombinant CCL3 added at 30 nM concentration promoted comparable numbers of migrating cells. Therefore, CCL3 activity appeared to be unaffected by the immobilization on the OMV surface. By contrast, the *in vitro* activity of immobilized TNFα and IL-2 was reduced by approximately 100-fold with respect to the soluble forms. The activity of the MBP-Flt3L fusion was also substantially lower than the purified recombinant Flt3L.

Overall, our data indicate that cytokines can be efficiently loaded on OMV membrane and they preserve at least part of their *in vitro* functional activity.

### Combinations of cytokine-bearing OMVs can induce complete tumor regression in ISV

Next, we tested whether the cytokine-expressing OMVs_Δ60_ had a superior *in vivo* anti-tumor activity with respect to the anti-tumor activity of the “Empty”-OMVs_Δ60_ (vesicles not carrying any cytokine). Inbred mice were challenged with different syngeneic tumor cell lines (colorectal cancer CT26 and MC38 cells, and WEHI-164 fibrosarcoma cells). Once tumors reached an average size of approximately 70-80 mm^3^, either engineered or “Empty” OMVs_Δ60_ (hereinafter defined as OMVs_Δ60_) were injected three times, two days apart, into the tumor mass and tumor growth was followed over a period of 60 or 80 days (Figure 3A and Figure 4A).

**Figure 3.**
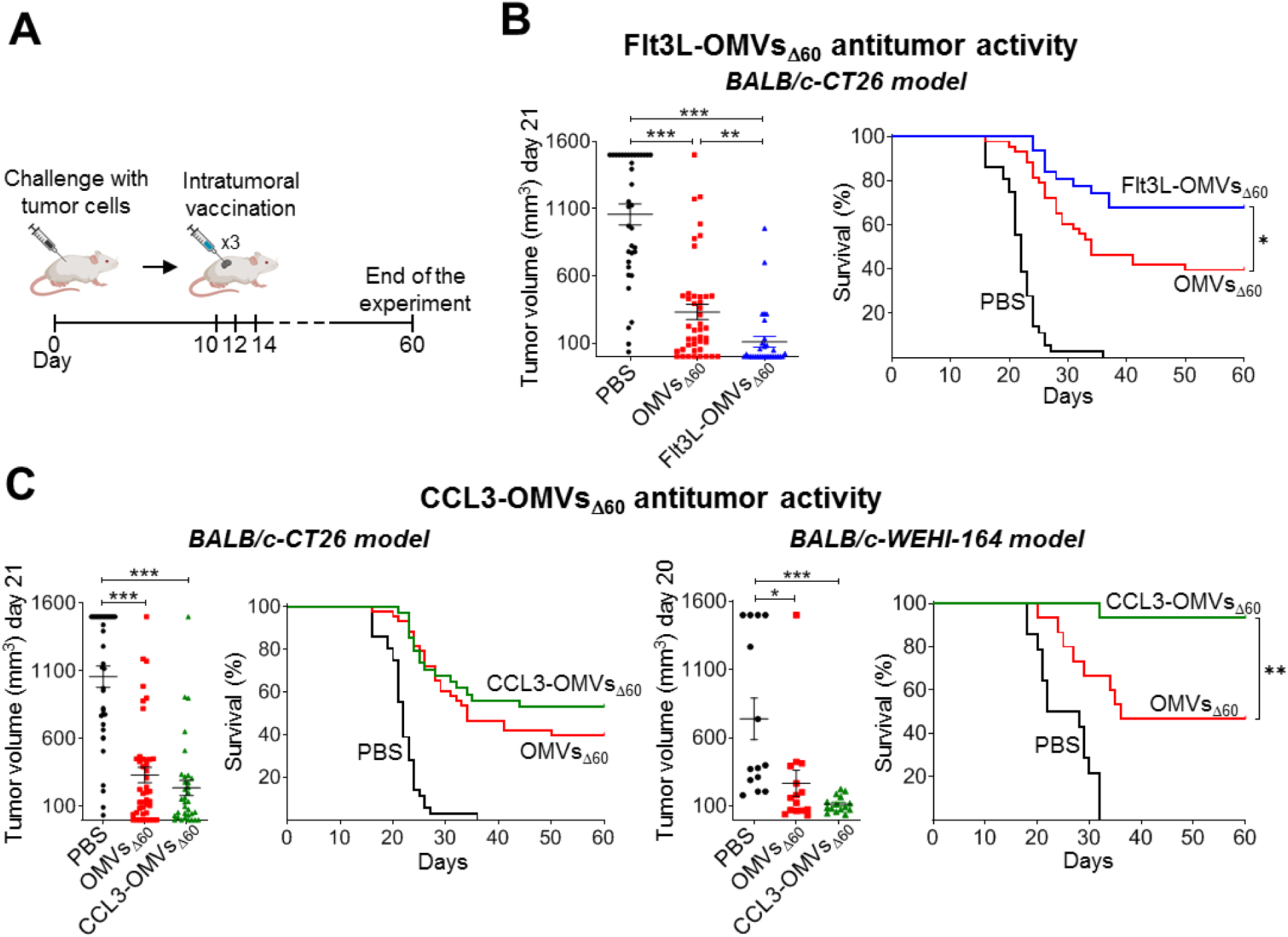
Tumor suppressing activity of CCL3-OMVs_Δ60_ and Flt3L-OMVs_Δ60_. (**A**) *Schematic representation of animal challenge and immunotherapy -* Eight-week-old female BALB/c mice were subcutaneously inoculated with syngeneic tumor cells and when tumors reached an average volume between 70-80 mm³, mice received three intratumoral injections two days apart of the appropriate formulations. Tumor size was monitored over a period of 60 days. (**B**) *Anti-tumor activity of Flt3L-OMV _Δ60_* – *BALB/c-CT26 model* - Mice were challenged with CT26 cells and tumors were treated with either PBS, or OMVs_Δ60_ (10 µg/dose), or Flt3L-OMVs_Δ60_ (10 µg/dose) (total number of animals treated in five independent experiments: PBS: 36 mice; OMV_Δ60_: 43 mice; Flt3L-OMV_Δ60_: 31 mice). Left graph shows the tumor size of all mice at day 21, while the right graph represents the Kaplan-Meier survival curves of each group. (**C**) *Anti-tumor activity of CCL3-OMV_Δ60_* – *BALB/c-CT26 model* – Animals were challenged with CT26 cells and tumors were treated with either PBS, or OMVs_Δ60_ (10 µg/dose), or CCL3-OMVs_Δ60_ (10 µg/dose) (total number of animals treated in five independent experiments: PBS: 36 mice; OMV_Δ60_: 43 mice; CCL3-OMV_Δ60_: 34 mice). Left graph shows the tumor size of all mice at day 21, while the right graph represents the Kaplan-Meier survival curves of each group. *BALB/c-WEHI-164 model* - Mice were challenged with WEHI-164 cells and tumors were treated as described above (total number of animals treated in three independent experiments: PBS: 14 mice; OMV_Δ60_: 15 mice; CCL3-OMV_Δ60_: 15 mice). Left graph shows the tumor size of all mice at day 20, while the right graph represents the Kaplan-Meier survival curves of each group. Statistical analysis of the Kaplan-Meier curves was performed using Gehan–Breslow–Wilcoxon Test while unpaired, two-tailed Student’s *t*-test was used for the statistical analysis of tumor volumes (* p ≤ 0.05; ** p ≤ 0.01; *** p ≤ 0.001).

**Figure 4.**
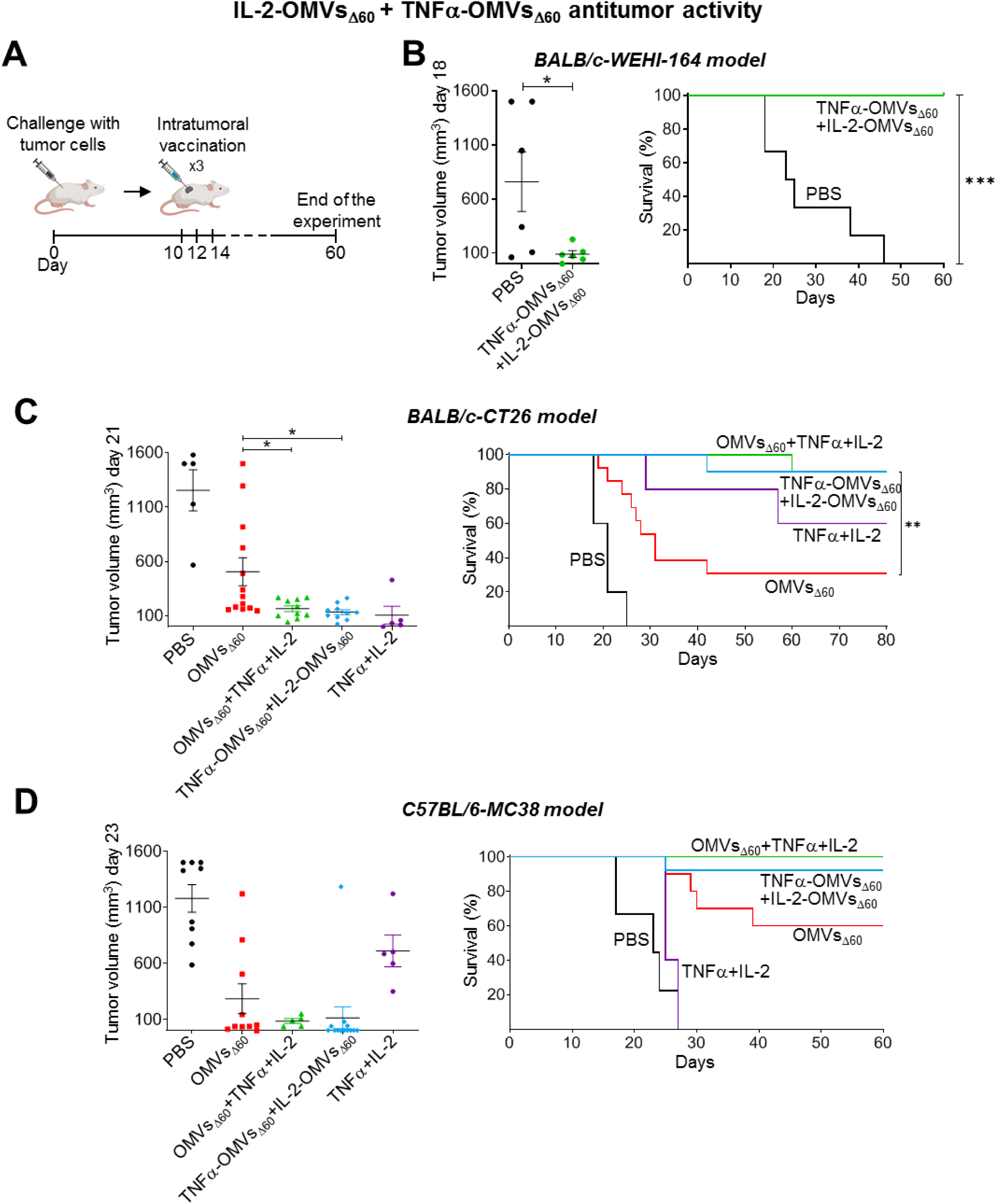
Tumor suppressing activity of TNFa-OMVs_Δ60_ + IL-2-OMVs_Δ60_ Combo. (**A**) *Schematic representation of animal challenge and immunotherapy -* Eight-week-old female BALB/c mice or C57BL/6 mice were subcutaneously inoculated with syngeneic tumor cells and when tumors reached an average volume between 70-80 mm³, mice received three intratumoral injections two days apart of the appropriate formulation. Tumor size was monitored over a period of 60 or 80 days. (**B**) *BALB/c-WEHI-164 model* - Mice (6 animals per group) were challenged with WEHI-164 cells and tumors were treated with either PBS or TNFα-OMVs_Δ60_ + IL-2-OMVs_Δ60_ Combo (5 µg of each engineered OMV/dose). Left graph shows the tumor size of all mice at day 18, while the right graph represents the Kaplan-Meier survival curves of each group. (**C**) *BALB/c-CT26 model* - Mice were challenged with CT26 cells and tumors were treated with either PBS, or OMVs_Δ60_ (10μg/dose), or OMVs_Δ60_ (10 μg/dose) + recombinant TNFα + recombinant IL-2 (150 ng each/dose), or TNFα-OMVs_Δ60_ + IL-2-OMVs_Δ60_ Combo (5 μg each/dose) (total number of mice per group: PBS: 5 mice; OMVs_Δ60_: 10 mice; OMVs_Δ60_ + TNF + IL-2: 10 mice; TNFα-OMVs_Δ60_ + IL-2-OMVs_Δ60_: 10 mice; TNFα + IL-2: 5 mice). Left graph shows the tumor size of all mice at day 21, while the right graph represents the Kaplan-Meier survival curves of each group. (**D**) *C57BL/6-MC38 model* - Mice were challenged with MC38 cells and tumors were treated as described in C) (total number of animals treated: PBS: 9 mice; OMVs_Δ60_: 10 mice; OMVs_Δ60_ + TNF + IL-2: 5 mice; TNFα-OMVs_Δ60_ + IL-2-OMVs_Δ60_: 13 mice; TNFα + IL-2: 5 mice). Left graph shows the tumor size of all mice at day 23, while the right graph represents the Kaplan-Meier survival curves of each group. Statistical analysis of the Kaplan-Meier curves was performed using Gehan–Breslow–Wilcoxon Test, while unpaired, two-tailed Student’s *t*-test was used for the statistical analysis of tumor volumes (* p ≤ 0.05; ** p ≤ 0.01; *** p ≤ 0.001).

In a first set of experiments, the tumor suppressing activity of Flt3L-OMVs_Δ60_ was tested using the BALB/c – CT26 mouse model. Three groups of animals were intratumorally injected, in five independent experiments, with either PBS, or OMVs_Δ60_, or Flt3L-OMVs_Δ60_. As shown in Figure 3B, 21 days after injection, tumor inhibition by Flt3L-OMVs_Δ60_ was improved in a statistically significant manner with respect to OMVs_Δ60_. Importantly, 67,7% of all Flt3L-OMVs_Δ60_-treated mice completely recovered from the tumor challenge as judged by the 60-day follow-up analysis. Such protection was statistically higher than the protection observed in OMVs_Δ60_-treated mice (39,5%) (Figure 3B).

Next, we tested the anti-tumor activity of CCL3-OMVs_Δ60_. Mice were intratumorally injected with CCL3-OMVs_Δ60_ and tumor growth was followed over a period of 60 days and compared to the tumor development in PBS- and OMVs_Δ60_–treated mice. As shown in Figure 3C (left panel), CCL3-OMVs_Δ60_ injection inhibited tumor growth and improved overall survival of CT26 injected mice more effectively than OMVs_Δ60_ (53% vs 39.5% - Figure 3C, left panels) although the difference did not reach statistical significance.

The CCL3-OMVs_Δ60_ anti-tumor activity was also tested in a second tumor mouse model. BALB/c mice were challenged with WEHI-164 fibrosarcoma cells and subsequently treated with either PBS or OMVs_Δ60_ or CCL3-OMVs_Δ60_. CCL3-OMVs_Δ60_ turned out to be remarkably protective in this tumor model, counteracting tumor growth and promoting complete tumor remission in 93.3% of treated mice, as opposed to 46.6% of OMVs_Δ60_ (Figure 3C, right panels).

A last set of experiments was carried out to test the antitumor activity of the combination of two engineered OMVs, one expressing TNFα (TNFα-OMVs_Δ60_) and the other expressing IL-2 (IL-2-OMVs_Δ60_). The selection of this OMV combo was based on the efficacy of these two cytokines (in the form of “immunocytokines”) for the treatment of skin cancer patients when administered intratumorally (*25, 26*).

In a first study, two groups of BALB/c mice (six mice per group) were challenged with WEHI-164 sarcoma cells and when tumors developed, one group was treated with PBS and the other with the TNFα-OMVs_Δ60_ + IL-2-OMVs_Δ60_ Combo (Figure 4A). As shown in Figure 4B, all mice treated with the OMV combo completely recovered from the tumor challenge.

Next, five groups of BALB/c mice bearing CT26 tumors (∼70–80 mm³) received three doses, two days apart (Figure 4A), of: (1) saline, (2) OMVs_Δ60_, (3) TNFα-OMVs_Δ60_ + IL-2-OMVs_Δ60_, (4) 150ng each of recombinant TNFα and IL-2 (amounts matched to cytokine-loaded OMVs), or (5) OMVs_Δ60_ + 150ng each of recombinant cytokines. Tumor growth was followed for 80 days. As shown in Figure 4C, OMVs_Δ60_ alone cured 39.5% of mice, whereas both cytokine-engineered OMVs_Δ60_ and OMVs_Δ60_ combined with recombinant cytokines achieved 90% complete responses. Recombinant TNFα + IL-2 alone delayed tumor growth in all mice and ultimately cured 60% of animals. Thus, TNFα and IL-2 synergise with OMVs_Δ60_ markedly enhancing anti-tumor efficacy of ISV in the CT26 model.

The study was repeated in the C57BL/6-MC38 mouse tumor model. Consistent with the BALB/c-CT26 results, OMVs_Δ60_ + recombinant cytokines and the TNFα-OMVs_Δ60_ + IL-2-OMVs_Δ60_ Combo cured 100% and 92.3% of mice, respectively (Figure 4D). In contrast, differently from what observed in the BALB/c-CT26 model, recombinant cytokines alone only transiently slowed tumor growth and all mice progressed by day 30.

In conclusion, our data strongly indicate that cytokines and OMVs act synergistically when co-delivered intratumorally. This synergy is most evident in the robust therapeutic activity of the TNFα-OMVs_Δ60_ + IL-2-OMVs_Δ60_ Combo across all mouse models tested.

### OMV administration modifies the tumor microenvironment

We previously showed that soon after intratumoral injection of OMVs_Δ60_, a strong inflammatory response develops, leading to the recruitment of NK cells and dendritic cells, together with extensive tumor-cell death and aggregation (*14*).

In a first set of experiments, we analyzed the effect on tumor microenvironment of TNFα-OMVs_Δ60_ + IL-2-OMVs_Δ60_ Combo, the formulation which completely inhibited tumor growth in all three models tested. Two tumors from BALB/c mice challenged with CT26 cells were collected 24 hours after three injections of either PBS or OMV Combo and the presence of myeloid cells and T cells in tumor sections were analyzed by immunostaining. As shown in Figure 5, differently from PBS injection, the ISV treatment caused a marked necrosis in the tumor mass, as reveled by the presence of extensive areas of amorphous tissue deprived of live cells. However, where live cells were present, they were surrounded by a rich infiltration of both CD8⁺ T cells and CD4⁺ T cells (Figure 5A-B). The number of total T cells in the section and of T cells per mm^2^ of section was substantially higher in OMV-treated tumors with respect to control tumors.

**Figure 5.**
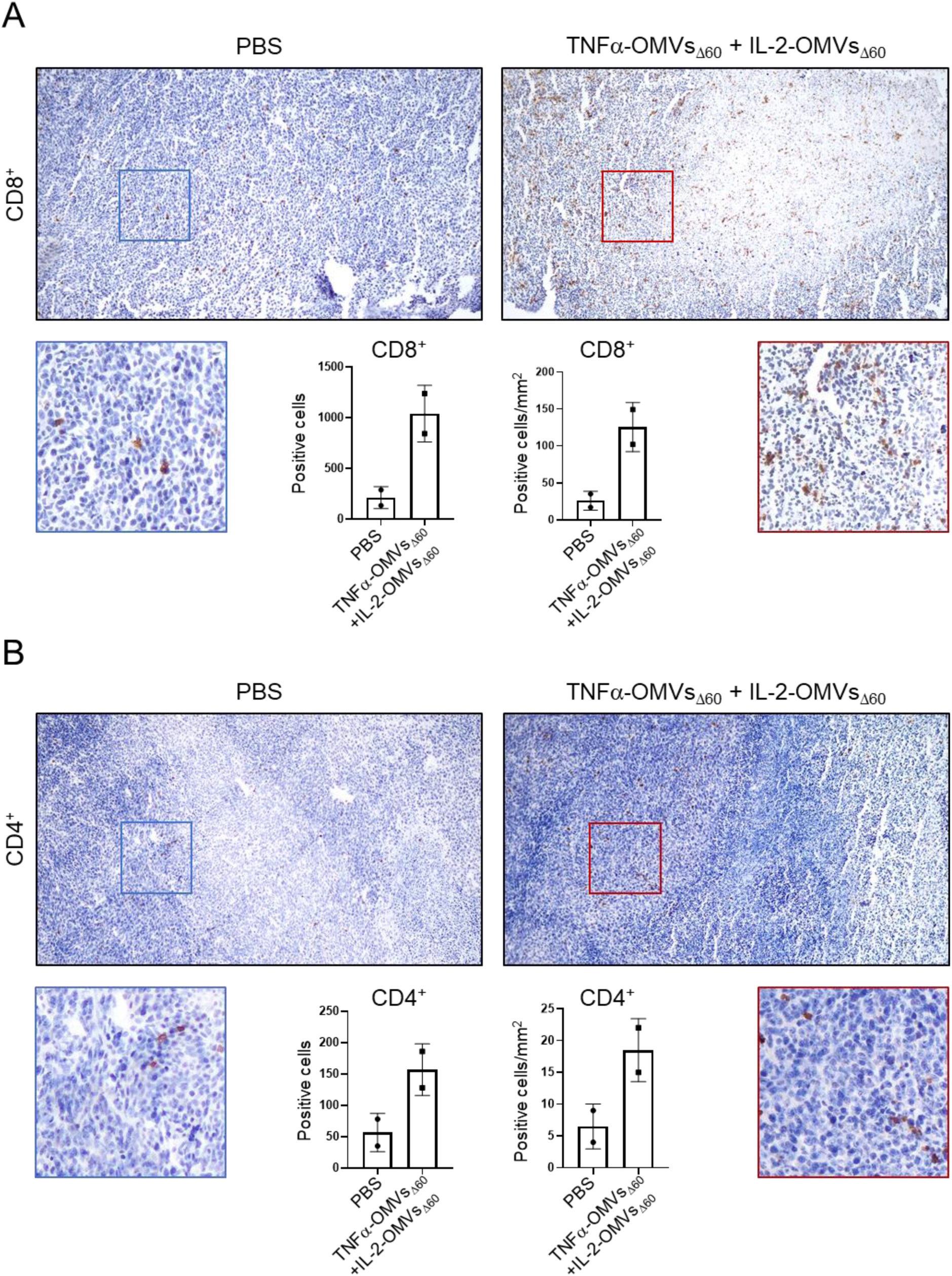
Immunohistochemistry analysis of CT26 tumors. CT26 tumors were collected 24 hours after a three intratumoral injections of PBS (50 µL) or TNFα-OMVs_Δ60_ + IL-2-OMVs_Δ60_ (10 µg in 50 µL PBS). Representative images of FFPE tumor sections stained with ematoxylin and for CD8^+^ (**A**) or CD4^+^ (**B**) T-cells. Bar plots represent the quantification of total positive cells and positive cells/mm^2^ with QuPath software (n=2 tumors injected with PBS; n=2 tumors injected with IL-2-OMVs_Δ60_ + TNFα-OMVs_Δ60_).

When tumor sections were stained with anti-MPO antibodies to visualize neutrophils and monocytes, MPO-positive cells were present in both the control and OMV-treated tumors. Interestingly, in OMV-treated tumors, MPO^+^ cells appeared to accumulate in large numbers at the border of the necrotic areas (Figure 6A-B).

**Figure 6.**
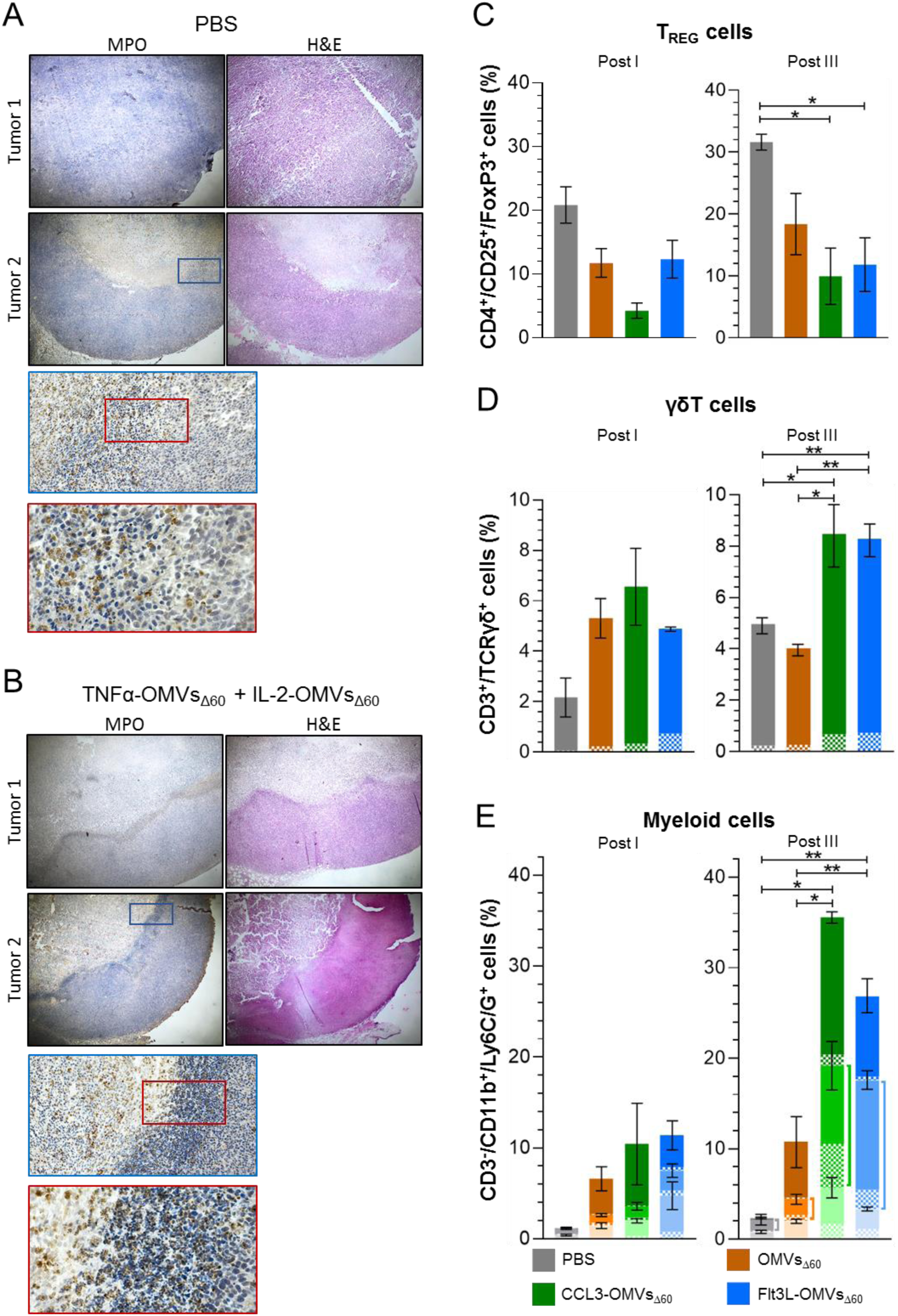
H&E-immunoistochemistry staining and flow cytometry analysis of CT26 tumors. Representative image of H&E staining MPO immunohistochemistry on FFPE sections of the same tumors described in Figure 5. CT26 tumors were collected 24 hours after a three intratumoral injections of PBS (50 µL) (**A**) or IL-2-OMVs_Δ60_ + TNFα-OMVs_Δ60_ (10 µg in 50 µL PBS) (**B**).(**C - D - E**) Flow cytometry analysis of tumors *–* BALB/c mice were challenged with CT26 and when tumors reached a size of approximately 100 mm^3^ mice were treated with one or three doses (two days apart) of the following formulations: PBS (control), 1 μg of OMVs_Δ60_, 1 μg of CCL3-OMVs_Δ60_, 1 μg of Flt3L-OMVs_Δ60_. The day after the treatments, two tumors from each group receiving one dose (Post I) and three tumors from each group receiving three doses (Post III) were surgically removed. Tumor cells (1 x 10^6^) were incubated with the appropriate fluorescent labelled antibodies and subsequently an alyzed by flow cytometry. Frequencies of regulatory T cells (***C***) γδ T cells (**D**) and LY6C/G^+^ CD11b^+^ myeloid cells (***E***) are calculated within the live, non-aggregated total cell populations. The dotted areas within the bars of ***panel D*** indicate the percentage of MHC II-positive γδ T cells. In ***panel E***, each bar is subdivided into three portions of different color intensity. The dark tone represents the fraction of LY6C/G^HIGH^CD11b^MEDIUM^ cells, the intermediate tone represents the fraction of LY6C/G^MEDIUM^ CD11b^HIGH^ cells and the light tone represents the fraction of myeloid cells excluded by the selected gating parameters. Finally, the dotted areas within each bar represent the fraction of each cell population which is MHC II-positive. The statistical analysis (unpaired, two-tailed Student’s *t*-test) shown in the right-hand graph refers to the populations of LY6C/G^MEDIUM^CD11b^HIGH^ cells present in each group (indicated by the lines next to the bars).

Overall, these data indicate that the therapeutic effect of TNFα-OMVs_Δ60_ + IL-2-OMVs_Δ60_ Combo is likely mediated by the concerted activation of both innate and adaptive immune responses.

Next, we also investigated the anti-tumor immune activity of CCL3-OMVs_Δ60_ and Flt3L-OMVs_Δ60_, in CT26 tumors harvested either 24 hours after a single treatment (Post I) or 24 hours after three injections (Post III). Tumor samples were analyzed by flow cytometry after sorting the live, single cell compartment (Figure S2). To minimize excessive cell death and aggregation a reduced dose of OMVs (1 μg/injection) was used.

CD3⁺CD4⁺ T cells and CD3⁺CD8⁺ T cells were detected within the viable single-cell fraction of OMV-treated tumors (Figure S2). The most striking effect of CCL3-OMVs_Δ60_ and Flt3L-OMVs_Δ60_ injections on the CD3^+^ population was observed in the of CD3⁺CD4⁺CD25⁺FoxP3⁺ subset (Regulatory T cells - Tregs). Indeed, the analysis of Tregs revealed a statistically significant reduction in their frequency in tumors treated with three doses of CCL3-OMVs_Δ60_ and Flt3L-OMVs_Δ60_ (Figure 6C and Figure S3). This is particularly interesting, considering that a low frequency of infiltrating Tregs is a hallmark of favorable prognosis in cancer patients (*30, 31*).

A second interesting immune cell population found in the OMV-treated tumors were the γδ T-cells. Our flow cytometry analysis revealed the presence of CD3⁺CD4⁻CD8⁻TCRαβ⁻ cells 24 hours after a single injection of OMVs, including OMVs_Δ60_. Staining with a γδ TCR-specific monoclonal antibody indicated that these cells belonged to the γδ T-cell subset. γδ T cells remained detectable after three OMV injections and were further enriched in tumors treated with OMVs carrying CCL3 and Flt3L (Figure 6D and Figure S4). γδ T cells are increasingly recognized as important contributors to antitumor immunity due to their potent MHC-independent cytotoxicity and their capacity to function as antigen-presenting cells (*32, 33*). Consistent with this, we found that approximately 8 to 10% of tumor-infiltrating γδ T cells in CCL3-OMV_Δ60_- and Flt3L-OMV_Δ60_-treated tumors expressed MHCII (Figure 6D and Figure S4).

The most striking alteration of the TME following OMV administration was the robust infiltration of CD3⁻CD11b⁺Ly6C⁺/Ly6G⁺ myeloid cells. These cells, representing inflammatory monocytes and neutrophils (the monoclonal antibody used recognizes a common epitope on Ly-6G and Ly-6C, previously known as the myeloid differentiation antigen Gr-1), accumulated in tumors within 24 hours from the first injection of both empty and engineered OMVs_Δ60_ and primarily had a phenotype CD3⁻CD11b^MED^Ly6C/G^HIGH^ (Figure 6E and Figure S5). After three injections, the total number of CD3⁻CD11b⁺Ly6C⁺/Ly6G⁺ cells markedly increased (Figure 6E and Figure S6), and these cells segregated into two prominent populations: CD3⁻CD11b^MED^Ly6C/G^HIGH^ and CD3⁻CD11b^HIGH^Ly6C/G^MED^. Notably, the CD3⁻CD11b^HIGH^Ly6C/G^MED^ population was particularly abundant in tumors treated with CCL3-OMVs_Δ60_ and Flt3L-OMVs_Δ60_. Finally, since it has been recently shown that Ly6C⁺ myeloid cells play an important role in antitumor immunity and, by expressing MHCII, can acquire antigen-presenting functions (*34*), we analyzed the presence of MHCII positive cells in our populations of CD3⁻CD11b⁺Ly6C⁺/Ly6G⁺ myeloid cells. We found that three OMV injections promoted the appearance of a considerable number of MHCII^+^ cells, and these cells were particularly abundant in the CD3⁻CD11b^HIGH^Ly6C/G^MED^ populations of both CCL3-OMV_Δ60_-treated tumors (45%) and Flt3L-OMV_Δ60_-treated tumors (35%) (Figure 6E and Figures S5 and S6).

## DISCUSSION

A defining feature of all cancer immunotherapy strategies is their shared objective to induce robust infiltration of immune cells from both myeloid and lymphoid lineages into the tumor microenvironment, ultimately leading to cancer cell elimination through diverse immunological mechanisms.

Among current immunotherapeutic approaches, intratumoral immunotherapy (ISV), which relies on the local delivery of inflammatory mediators and/or tumor-lytic agents, has emerged as a particularly promising strategy due to its simplicity, limited toxicity, and broad applicability. Based on the assumption that the tumor microenvironment already harbors a diverse repertoire of tumor antigens, ISV—unlike personalized vaccines—does not require prior identification of antigenic targets. The main goal of ISV is to activate immune mechanisms within the tumor microenvironment to promote recognition of tumor antigens and the induction of tumor-specific effector B and T cells. Once locally activated, these lymphocytes can disseminate systemically and potentially target distant metastases. Furthermore, because the immunostimulatory agents are administered directly into the tumor, lower doses are sufficient to achieve therapeutic efficacy, thereby minimizing systemic toxicity and immune-related adverse effects.

The efficacy of ISV critically depends on the formulation injected into the tumor. An optimal formulation should attract dendritic cells specialized in antigen cross-presentation and antitumor myeloid populations, facilitate tumor antigen spreading, and promote the expansion of tumor-specific effector T and B cells.

Here, we report a novel formulation particularly suited for ISV. Leveraging the intrinsic properties of OMVs, which activate innate immunity through microbe-associated molecular patterns (MAMPs) and induce immunogenic cell death via pyroptosis, we engineered OMVs to incorporate selected cytokines. This design integrates MAMP-driven inflammation of the tumor microenvironment, OMV-mediated dissemination of damaged-associated molecular patterns (DAMPs) and tumor antigens, and cytokine-dependent immune cell recruitment and activation into a single platform. Together, these mechanisms elicit complete tumor regression across all murine models we tested to date.

A first notable finding of our study is the efficiency with which cytokines and chemokines can be expressed in OMVs. Using the staphylococcal lipoprotein FhuD2 as a carrier, we successfully delivered a broad panel of cytokines from different species into the vesicular compartment. Importantly, confocal microscopy of engineered strains and dot blot analysis of OMVs indicated that these proteins protrude from the membrane surface. Surface exposure is likely to represent a critical feature of our platform, as cytokines must engage their cognate receptors to exert their biological functions, an interaction that could be hindered if the cytokines were confined within the vesicle lumen.

Regarding cytokine functionality, another key result of this study is that OMV-immobilized cytokines retain measurable *in vitro* activity. This is at least true for CCL3, TNFα and IL-2, three of the four cytokines we focused our attention on. However, except CCL3, which maintains a strong capacity to recruit PBMCs, both TNFα and IL-2 (and likely Flt3L as well) display reduced activity. This observation is not unexpected, given that cytokines are naturally secreted molecules that normally diffuse through the extracellular milieu to reach and activate their target cells.

Perhaps the most striking finding of our work is that, despite the reduced *in vitro* functional activity of OMVs_Δ60_-associated cytokines, the *in vivo* antitumor response of OMVs_Δ60_ is markedly enhanced when the vesicles are decorated with cytokines. We observed this enhancing effect with CCL3-OMVs_Δ60_ and Flt3L-OMVs_Δ60_, the injection of which resulted in a statistically significant increase of the OMVs_Δ60_ therapeutic index in at least one tumor mouse model. However, the best results were obtained with TNFα-OMVs_Δ60_ and IL-2-OMVs_Δ60_ Combo, which eradicated tumors in almost 100% of mice challenged with three different syngeneic cancer cell lines. In these experiments the synergistic action of OMVs_Δ60_ and cytokines clearly emerges. This is particularly evident in the C57BL/6-MC38 model where the intratumoral co-injection of purified recombinant TNFα+IL-2 does not affect the overall animal survival, while full protection is observed when the cytokines are either mixed with OMVs_Δ60_ or engineered in the vesicles. It is interesting to note that despite the common origin of MC38 and CT26 tumors, while the administration of recombinant TNFα+IL-2 formulation alone was not effective against MC38 tumors, it conferred approximately 60% protection in the BALB/c-CT26 model. This is in line with the reported immunosuppressive TME of MC38 allografts (*43*) and, accordingly, to the higher efficacy of ICI treatment in BALB/c-CT26 model with respect to the C57BL/6-MC38 model (*44*). The synergistic action of OMVs_Δ60_ and cytokines strongly suggests that cytokine presentation within the context of OMVs provides additional immunostimulatory cues or localization advantages that also compensate for the reduced intrinsic *in vitro* activity of cytokines immobilized on the OMV surface. This also opens the possibility of a combined approach using ICIs and OMVs in immune refractory tumors.

In this study, to gain insight into the immunological mechanisms underlying the antitumor activity of OMVs, we analyzed the tumor microenvironment (TME) using immunohistochemistry (IHC) of formalin-fixed paraffin-embedded (FFPE) tumor sections and flow cytometry.

From the IHC analysis of tumors treated with TNFα-OMVs_Δ60_ + IL-2-OMVs_Δ60_ Combo it clearly emerged that after three injections extensive necrosis within the tumor mass occurred, accompanied by a massive recruitment of myeloid cells (neutrophils and monocytes) and CD4^+^ and CD8^+^ T cells, strongly indicating that the potent therapeutic efficacy of the OMV Combo is mediated by the concerted activation of both innate and adaptive immune responses. Interestingly, the infiltration of T cells in tumors is in line with the data previously published by Danielli and colleagues, showing that in melanoma patients the anti-tumor activity of the L19-IL2/L19-TNFα injection resulted in a potent CD4^+^/CD8^+^ T cell infiltration [25].

Regarding the flow cytometry analysis of tumors treated with CCL3-OMVs and with Flt3L-OMVs, a few noteworthy observations emerged.

First, consistent with the inflammatory properties of OMVs and their capacity to activate innate immunity, we found that CD11b⁺Ly6C/G⁺ myeloid cells were recruited to the tumor site 24 hours after a single OMVs_Δ60_ administration. This population appeared to be more abundant in tumors treated with CCL3-OMVs_Δ60_ and Flt3L-OMVs_Δ60_. Notably, after three injections of both empty and engineered OMVs_Δ60_, not only did the frequency of CD11b⁺Ly6C/G⁺ cells further increase, but these cells also segregated into two distinct populations, one of which contained a substantial fraction of MHCII⁺ cells. Such MHCII^+^ population was particularly abundant in tumors treated with CCL3-OMVs_Δ60_ and Flt3L-OMVs_Δ60_ (Figure 6). The OMV-mediated accumulation of MHCII⁺ myeloid cells in tumors resembles what happened in cancer patients who received COVID-19 mRNA vaccination. Grippin and colleagues reported that immune-checkpoint inhibitor efficacy was significantly enhanced in patients vaccinated within 100 days of treatment initiation (*34*). They demonstrated that the beneficial effect of vaccination on overall patient survival was mediated by the MDA5- dependent activation of myeloid cells—particularly Ly6C⁺ MHCII⁺ cells—which accumulated in tumors and promoted antigen presentation and the expansion of tumor-specific T cells.

Because of the extensive inflammation occurring soon after treatment, we did not perform flow cytometry on tumors receiving TNFα-OMVs_Δ60_ + IL-2-OMVs_Δ60_ Combo. However, considering that “Empty” OMVs_Δ60_ administration was sufficient to promote the recruitment of myeloid cells, we predict that a similar effect is likely elicited by the cytokine-bearing-OMV combination treatment.

A second interesting observation emerging from our study is the presence of γδ T cells in OMV-treated tumors. We found a rapid appearance of γδ T cells in the tumor microenvironment following a single OMVs_Δ60_ injection, with their frequency increasing further after three administrations of CCL3-OMVs_Δ60_ or Flt3L-OMVs_Δ60_. An appreciable proportion of these cells (approximately 10%) displayed an activated phenotype, particularly in Flt3L- OMVs_Δ60_ treated tumors, as indicated by MHCII expression. γδ T cells are increasingly recognized as potent mediators of antitumor immunity, capable of detecting and eliminating stressed or transformed cells through both TCR- and NKG2D-dependent mechanisms (*35–37*). In particular, the Vγ9Vδ2 TCR recognizes phosphoantigens (pAgs) that accumulate in tumor cells, while the NKG2D receptor binds MICA/MICB molecules often overexpressed on cancer cells. Interestingly, γδ TCRs also recognize microbial metabolites such as (E)-4-hydroxy-3-methyl-but-2-enyl pyrophosphate (HMBPP), produced by bacteria including *E. coli* (*38, 39*). Therefore, it is plausible that OMVs derived from *E. coli* contain sufficient HMBPP to directly activate γδ T cells. Upon activation, γδ T cells exert cytotoxic activity via perforin, granzymes, and TRAIL, mediate antibody-dependent cytotoxicity through CD16, secrete IFNγ and other Th1 cytokines, and can cross-present antigens to αβ T cells, thus amplifying adaptive responses (*35, 37*). Taking together, these findings suggest that the tumor-suppressive effects of intratumoral OMVs can be in part mediated by γδ T-cell activation and infiltration, particularly when vesicles are engineered with cytokines such as Flt3L. Since IL-2 serves as a potent proliferative and survival factor for γδ T cells following initial activation through TCR or NKG2D engagement, it is also plausible to believe that, similarly to what happens in tumors treated with CCL3-OMVs_Δ60_ and Flt3L-OMVs_Δ60_, the TNFα-OMVs_Δ60_ + IL2-OMVs_Δ60_ Combo promotes γδ T-cell–mediated antitumor responses.

A third relevant observation is related to the T cell compartment. Our immunostaining of OMV-treated tumors revealed that the non-necrotic areas were densely populated by CD8^+^ T cells and CD4^+^ Th1 T cells and, consistently with the T cell recruiting and/or activation capacity of cytokines, this was particularly evident in tumors treated with cytokine-bearing OMVs. Moreover, our flow cytometry data show that while tumors from control animals were invaded by a considerable population CD4^+^CD25^+^FoxP3^+^ T cells (Tregs) (approximately 20% and 30% of total CD4^+^ T cells after one and three PBS injections, respectively), the frequency of Treg cells dropped down by two-/three-fold in OMVs treated tumors and this was true for both OMVs_Δ60_ and engineered OMVs_Δ60_. The importance of αβ T cell populations in cancer immunity is extensively documented. The abundance of tumor-infiltrating cytotoxic CD8^+^ T cells and Th1 CD4^+^ T cells positively correlates with favorable prognosis, while CD4^+^FoxP3^+^ regulatory T cells inversely correlate with overall survival and therapeutic efficacy (*30, 31*).

Altogether, the results of our analysis demonstrate that OMV administration profoundly remodels TME, driving the recruitment of inflammatory myeloid cells, the recruitment of γδ T cells, the accumulation of αβ CD8^+^ and CD4^+^ T cells, and the depletion of Tregs. The convergence of these effects can well explain the potent therapeutic activity of cytokine-bearing OMVs and underscores their promise as a versatile platform for intratumoral immunotherapy.

Based on our preclinical results, the TNFα-OMVs_Δ60_ + IL-2-OMVs_Δ60_ Combo deserves testing in human patients. A first attractive target population are skin tumor patients, and particularly, patients affected by Basal Cell Carcinoma (BCC) and cutaneous Squamous Cell Carcinoma (cSCC). BCC and cSCC belong to keratinocyte carcinomas, the most common malignancies worldwide, with BCC accounting for about 75% of all keratinocyte cancers. The prognosis for cSCC patients and particularly for BCC patients is usually favorable, with a mortality rate ranging from 1.5% to 0.1%, respectively. The recommended treatment modality is surgery, but other treatments are available, such as topical therapy with imiquimod or 5-fluorouracil, photodynamic therapy and radiotherapy (*45*). However, for locally advanced, recurrent and metastatic BCC and for cSCC, whenever surgery is often no longer feasible, first- and second-line therapies are recommended and include radiotherapy, inhibitors of the Hedgehog pathway (Hh) and checkpoint inhibitors (anti-PD-1 antibodies). These therapies are only partially efficacious and very expensive (>100.000 Euros/patient). Therefore, new therapeutic strategies are urgently needed and indeed several clinical trials are in progress (https://clincaltrials.gov). Interestingly, intralesional treatments are a strong focus of ongoing clinical trials. The major limitation of our study is that the immunotherapeutic activity of our engineered OMVs is based on syngeneic mouse tumor models which do not fully recapitulate the biology and pathogenesis of cancer in humans. In the absence of reliable alternative mouse models, to mitigate this limitation, we tested the potency of our formulation against mouse tumors of different origin, differing in their genetic make-up, tumorigenic programs and TME. The fact that the TNFα-OMVs_Δ60_ + IL-2-OMVs_Δ60_ Combo consistently promotes tumor regression in three tumor models and similar results are being accumulated in other models, makes us confident of the therapeutic value of our approach.

While at present the TNFα-OMVs_Δ60_ + IL-2-OMVs_Δ60_ Combo is our most promising vaccine candidate, in this work we also demonstrated that several cytokines and chemokines other than CCL3, Flt3L, TNFα and IL-2 can be efficiently incorporated into the OMVs. Since many of them play relevant roles in tumor immunity (*16, 17, 19*) it would be interesting to extend the *in vivo* studies using different engineered OMVs and different OMV combinations. Considering the ease and the efficiency with which our engineered OMVs can be produced, such studies would help shed light on the cooperative action of cytokines in tumor immunity and might lead to the identification of optimized versions of OMV formulations tailored for the treatment of specific solid tumors in combination with checkpoint inhibitors and/or other conventional therapies and immunotherapy strategies. In this regard, we previously showed that the intratumoral injection of OMVs combined with cancer neoepitopes promotes a potent “abscopal effect,” as demonstrated in a two-tumor mouse model (*14*). We predict that this long-distance, immune-mediated anti-tumor activity would be further enhanced if neoepitopes were combined with our cytokine-bearing OMVs, thereby expanding their therapeutic potential to highly metastatic tumors.

## MATERIALS AND METHODS

### Cloning strategies

Cytokines were expressed in the OMVs by fusing them at the C-terminal of the *Staphylococcus aureus* FhuD2 lipoprotein. To do so, the synthetic genes encoding for murine cytokines (GeneArt, Thermo Fisher Scientific, Waltham, MA, USA) were PCR amplified (Q5 polymerase, New England Biolabs, Ipswich, MA, USA) adding the restriction sites for BamHI and XhoI at their 5’- and 3’-ends, respectively. PCR products were then digested for 1h at 37°C with BamHI and XhoI (New England Biolabs, Ipswich, MA, USA) and inserted into the vector pET21b-FhuD2, previously linearized by digestion with the same restriction enzymes, thus fusing the synthetic genes in frame to the 3’-end of FhuD2 coding sequence (the BamHI site sequence inserted a Glycine-Serine (GS) linker between the protein carrier and the cytokine). Ligation products were used to transform DH5α cells and positive clones were selected by colony PCR with GoTaq (Promega, Madison, WI, USA). The sequence of each construct was confirmed by DNA sequencing. Finally, the plasmids were used to transform the hypervesiculating *E. coli* BL21(DE3)Δ60 strain for OMVs production.

### OMVs preparation and purification

OMVs were prepared as described in Tamburini *et al*. (*40*). Briefly, overnight cultures (50 mL) were added to 450 mL of Luria-Bertani (LB) (Sigma-Aldrich, St. Louis, MO, USA) supplemented with 100 μg/mL ampicillin (Carl Roth GmbH & Co. KG, Germany) and bacterial growths were carried out at 30°C under shaking (200 rpm). At OD_600_ between 0.5 and 0.7, the expression of cytokine fusions was induced by adding isopropyl-β-D-thiogalactoside (IPTG, Carl Roth GmbH & Co. KG, Germany) at a final concentration of 0.1 mM. After 2-3 hours at the end of the exponential phase, the biomass of the cultures was separated from the supernatant by centrifugation at 6.000 × g at 4°C for 20 minutes. The supernatants were filtered through a 0.22 μm pore size filter (Merck, Darmstadt, Germany) and treated with 1 U/ml Benzonase® nuclease (Merck, Darmstadt, Germany) at 4°C for 15 hours. OMVs were purified from the supernatants using Tangential Flow Filtration (TFF) with a Cytiva Äkta Flux system (Cytiva, Marlborough, MA, USA) equipped with a Hollow Fibre cartridge UFP-500-C-3MA (Cytiva, Marlborough, MA, USA). After extensive dialysis with sterile PBS, OMVs were concentrated to reach a final OMV concentration of 1 mg/ml based on total protein content determined using the DC^TM^ Protein Assay Kit II (BioRad, Hercules, CA, USA). For protein profiling, OMV samples were resolved on Any kD Criterion TGX Stain-Free gels (Bio-Rad, Hercules, CA, USA) by electrophoresis and proteins were visualized by Coomassie Blue staining (Giotto, Sesto Fiorentino, Italy).

### 2D gel analysis of OMVs

Three technical replicates per condition, IL-2-OMVs_Δ60_ and TNFα-OMVs_Δ60_, were included for a differential proteomic analysis. OMVs were lyzed and proteins extracted by using a buffer consisting of Urea 7 M, Thiourea 2 M, 4% (w/v) 3-[(3-Cholamidopropyl)dimethylammonium]-1-propane sulfonate (CHAPS), 1,4-Dithioerythritol (DTE) 65 mM and 2% (*v*/*v*) Triton X-100. 250ug per each technical replicate were used to perform two-dimensional gel electrophoresis (2DE), in order to separate intact proteins according to their isoelectric point (pI) and molecular weight (MW), allowing for the discrimination of proteoforms. First, Isoelectric Focusing (IEF) was performed on 18 cm immobilized nonlinear pH 3–10 gradient IPG strips (Cytiva, Uppsala, Sweden; formerly GE Healthcare) by rehydration loading protocol using an Ettan™ IPGphor™ system (GE Healthcare, Uppsala, Sweden). Specifically, samples were supplemented with 0.2% carrier ampholytes and IPG strips were, first, rehydrated for 1 h at 0 V and then overnight at 30 V at 16°C. Following IEF steps included a specific voltage program: 200 V for 8 h, gradient from 200 V to 3500 V in 2 h, held at 3500 V for 2 h, gradient from 3500 V to 5000 V in 2 h, held at 5000 V for 3 h, gradient to 8000 V in 1 h, and maintained at 8000 V until reaching a total of 90,000 VhT. After IEF, IPG strips were rinsed with deionized water and equilibrated in a buffer containing 6 M urea, 2% *w*/*v* SDS, 2% *w*/*v* DTE, 30% *v*/*v* glycerol, and 0.05 M Tris-HCl pH 6.8 for 12 min, then incubated for 5 min in a similar buffer where DTE was replaced with 2.5% *w*/*v* iodoacetamide, with further addition of traces of bromophenol blue. Second dimension consisted in SDS-PAGE, carried out on 9–16% SDS–polyacrylamide linear gradient gels (18 × 20 cm, 1.5 mm thick), applying constant current of 40 mA per gel at 9 °C. 2DE gels were stained with ammoniacal silver staining and then digitalized using an Image Scanner III laser densitometer equipped with LabScan 6.0 software (GE Healthcare). Image visualization and comparative analysis were carried out using dedicated software Melanie™ Classic 9 (SIB Swiss Institute of Bioinformatics, Geneva, Switzerland). Comparative statistical analysis was carried out by ANOVA test, taking into account the percentage of relative volume (%V) of each spot and a p-value ≤ 0.05 and a fold change ≥ 4.5. Principal Component Analysis (PCA) by Spearman’s correlation and heatmap analysis by Euclidean distance were carried out using %V of differential protein species using XLStat (Addinsoft, Paris, France).

For 2DE immunoblots, 120 µg per condition, IL-2 OMVs and TNF-α OMVs, were used. After two-dimensional gel electrophoresis, 2DE gels were equilibrated for 1 h in Towbin transfer buffer (Tris 25 mM, glycine 192 mM, methanol 20% (v/v)) and then, transferred onto nitrocellulose membrane (Amersham Hybond ECL Nitrocellulose Membrane, Cytiva, Marlborough, MA, USA). Membranes were stained by Red Ponceau (ponceau S 0.2% (w/v) in acetic acid 3% (v/v)) in order to check correct transfer. Immunoblotting was performed as follows: 3 washes in blocking solution (powdered milk 3% (w/v), Triton X-100 0.1% (v/v) in PBS pH 7.4) for 10 minutes each, then incubation overnight at 4°C with primary antibody (diluted 1:1000); 3 washes in blocking solution for 10 minutes each, then incubation for 2 hours at room temperature with secondary antibody; 3 washes in blocking solution for 10 minutes each, then incubation for 30 minutes in Triton X-100 0.5% (v/v) in PBS pH 7.4 and 2 washing steps in 0.05 M Tris-HCl pH 6.8 for 30 minutes each. Immunoreaction was carried out by ECL chemiluminescence detection system and signals were detected by exposing membranes to Hyperfilm ECL X-ray films (Cytiva). 2DE-WB images were analysed by Melanie™ Classic 9 (SIB). Primary antibodies are rat anti-IL-2 (BioLegend, San Diego, CA, USA) and goat anti-TNFα (R&D Systems, Minneapolis, MN, USA). Specific HRP-conjugated secondary antibodies are rabbit anti-goat IgG (R&D Systems, Minneapolis, MN, USA), goat anti-rat IgG (Invitrogen, Waltham, MA, USA), diluted 1:1000.

### Confocal microscopy

Recombinant *E. coli* strains were grown in LB and protein expression induced at OD₆₀₀ = 0.4/0.6 with 0.1 mM IPTG as previously described. When cultures reached an OD₆₀₀ = 1, 1 mL of each culture was harvested by centrifugation at 6.000 x g for 2 minutes, bacteria were washed with PBS and fixed with 500 μl of Cytofix (BD Bioscience, San Jose, CA, USA) for 15 minutes at room temperature. Fixed cells were pelleted at 3.500 x g for 2 minutes, washed three times with PBS and resuspended in 1 mL of PBS. For staining, aliquots of fixed bacteria were processed under non-permeabilizing conditions. Briefly, after blocking with PBS 1% BSA, cells were stained with the following primary antibodies diluted at a final concentration of 1 µg/mL in PBS: anti-TNFα (R&D Systems, Minneapolis, MN, USA); anti-CCL3 (R&D Systems, Minneapolis, MN, USA); anti-IL-2 (BioLegend, San Diego, CA, USA); anti-Flt3L (R&D Systems, Minneapolis, MN, USA). After 1h incubation at RT in the dark, bacteria were washed three times with PBS 1% BSA and resuspended in the appropriate secondary antibody conjugated with Alexa Fluor 488 or 647 at a final dilution of 1:400 in PBS. For genomic DNA staining and bacteria visualization, DAPI at a final dilution of 1:5.000 was included during secondary antibody incubation. This secondary step was carried out at room temperature for 30 minutes in the dark. Finally, after three washes with PBS, samples were mounted on microscopy slides using ProLong™ Gold mounting medium (Thermo Fisher Scientific, Waltham, MA, USA) and examined by Nikon N-SIM confocal microscopy (Nikon Europe B.V., Amstelveen, The Netherlands).

### Dot-blot Assay

Flt3L-OMVs_Δ60_, CCL3-OMVs_Δ60_, IL-2-OMVs_Δ60_, TNFα-OMVs_Δ60_ and OMVs_Δ60_ (negative control) were spotted in duplicate onto a nitrocellulose membrane (2 μg/spot) and allowed to air-dry. Then the membranes were blocked for 1 hour at RT in 10% skimmed milk (Merck, Darmstadt, Germany) in TBPS (0.05% Tween 20 in PBS, pH 7.4) under mild agitation. Subsequently, the membranes were incubated for 1 hour at RT with the cytokine-specific primary antibody diluted at the appropriate concentration in 1% skimmed milk in TPBS. As a positive control, the primary antibody against the FhuD2 carrier protein was also included in the analysis. In detail, the following primary antibodies were used: goat anti-Flt3L (5µg/mL, R&D Systems, Minneapolis, MN, USA), goat anti-CCL3 (1µg/mL, R&D Systems, Minneapolis, MN, USA), rat anti-IL-2 (5µg/mL, BioLegend, San Diego, CA, USA), goat anti-TNFα (1µg/mL, R&D Systems, Minneapolis, MN, USA), and rabbit anti-FhuD2 (5µg/mL) (*28*). After three washing steps in TPBS, the filters were incubated for 1 hour with the specific HRP-conjugated secondary antibodies (rabbit anti-goat IgG (R&D Systems, Minneapolis, MN, USA), goat anti-rat IgG (Invitrogen, Waltham, MA, USA) and goat anti-rabbit IgG (Invitrogen, Waltham, MA, USA)) diluted 1:5.000 in 1% skimmed milk in TPBS. The membranes were washed three times with TPBS and after addition of the SuperSignal West Pico chemiluminescent substrate (Thermo Fisher Scientific, Waltham, MA, USA) the immunoreactive signals were detected with the ImageQuant LAS4000 (GE, Chicago, IL, USA) instrument.

### Determination of CCL3 in vitro activity: migration assay

Mouse PBMCs were isolated from the blood of four CD1 mice using the Ficoll gradient method. Blood was diluted 1:1 with PBS and carefully layered over Ficoll (1:2 Ficoll:blood). Samples were centrifuged at 400 × g for 30 minutes without brakes. The PBMC layer was collected, washed with 10 mL of RPMI medium, and centrifuged at 300 × g for 10 minutes. The pellet was resuspended in fresh RPMI. A total of 1.5 × 10⁵ PBMCs were seeded in the upper chamber of a Corning® 6.5 mm Transwell® with a 5.0 µm polycarbonate membrane. The lower chamber contained RPMI alone or supplemented with OMVs_Δ60_ (1 µg), CCL3-OMVs_Δ60_ (1 µg), or recombinant CCL3 (30 nM, R&D Systems, Minneapolis, MN, USA). Migrated cells in the lower chamber were counted using a Neubauer chamber. Graphs represent the percentage of input cells that migrated. Statistical analysis was performed using an unpaired *t*-test.

### Determination of IL-2 in vitro activity: proliferation assay

The biological activity of the OMV-associated cytokine IL-2 was determined by its ability to stimulate the proliferation of murine cytotoxic T lymphocytes (CTLL-2) (ATCC, Manassas, VA, USA) as described in Corbellari *et al*. (*41*). In 96-well plates, 2.5 × 10⁴ CTLL-2 cells in starvation (washed 24 hours before removal of the Rat-STIM supplement (BD Bioscience, San Jose, CA, USA)) were seeded per well in RPMI culture medium and then serial dilutions of either OMVs_Δ60_ or IL-2-OMVs_Δ60_ were added. Recombinant mouse IL-2 (BioLegend, San Diego, CA, USA) was used as a positive control. After incubation at 37°C for 72 hours, cell proliferation was determined using the CellTiter Aqueous One Solution® (Promega, Madison, WI, USA). Results were read at 490 nm – 620 nm after 4 hours and expressed as the percentage of cell viability compared with untreated cells.

### Determination of TNFα in vitro activity: cytotoxicity assay

For the cytotoxicity assay, MC38 carcinoma cells (2 × 10⁴ cells per well) were seeded in RPMI medium supplemented with 2 μg/mL actinomycin D (Sigma-Aldrich, St. Louis, MO, USA) and incubated with serial dilutions of either OMVs_Δ60_ or TNFα-OMVs_Δ60_ or recombinant mouse TNF (BioLegend, San Diego, CA, USA). After 24 hours, cytotoxicity was assessed using the CellTiter Aqueous One Solution® (Promega, Madison, WI, USA). Results were read at 490 nm - 620 nm after 3 hours and expressed as the percentage of cell viability compared with the control group (cells treated with actinomycin D only).

### Determination of Flt3L in vitro activity: proliferation assay

The biological activity of recombinant Flt3L and Flt3L fused to the Maltose-Binding Protein (MBP) was determined by following the proliferation of human acute myeloid leukemia OCI AML-5 cell line. The MBP-Flt3L fusion was generating adding the murine Flt3L gene in-frame to the 3′ end of the MBP gene using the strategy described in *Grandi A. et al.* (*42*). The generated pET21-MBP-Flt3L plasmid was used to transform *E. coli* BL21(DE3)Δ60. More specifically, the *E. coli* BL21(DE3)Δ60-(pET-MBP-Flt3L) recombinant strain was grown in LB to an OD₆₀₀ = 0.4/0.6, and then protein expression was induced by adding 0.1 mM IPTG. After two hours, the bacterial pellet was collected by centrifugation at 6.000 x g for 20 min. This pellet was lysed with B-PER™ Bacterial Protein Extraction Reagent (Thermo Fisher Scientific, Waltham, MA, USA) supplemented with 100 μg/mL Lysozyme, 1U/ml DNAse and 1mM MgCl_2_ for 30 minutes on ice. The lysate was then centrifuged at 11.000 x g for 10 min and the supernatant (soluble fraction) containing the MBP-Flt3L protein, collected. This soluble fraction was loaded into a MBPTrap HP 1mL column (Cytiva, Marlborough, MA, USA) previously equilibrated with binding buffer (20mM Tris-HCl, 200mM NaCl pH 7.6). After washing with 10 volumes of the column with binding buffer, the protein was eluted and collected with elution buffer (Binding buffer supplemented with 10mM Maltose). The quality of the protein was verified by SDS-PAGE and protein concentration was measured with the Bradford Protein Assay (BioRad, Hercules, CA, USA).

For the proliferation assay, OCI AML-5 cells were grown in Alfa-MEM medium supplemented with 20% of Fetal Bovine Serum (FBS) with the addition of 10 ng/mL of Granulocyte-macrophage colony-stimulating factor (GM-CSF). Before the assay, the cells were maintained for 48 hours in Alfa-MEM medium w/o GM-CSF and then plated in a 96-well plate at a concentration of 1.2 x 10^5^ cells/well. The cells were then subjected to different stimulations for 48 hours at 37°C. Specifically, cells were stimulated with different concentrations of recombinant Flt3L protein (Miltenyi Biotech, Bergisch Gladbach, Germany) as a positive control, and of MBP-Flt3L fusion recombinant protein. As negative control, an irrelevant MBP fusion proitein (MBP-Pfept), constituted by a *P. falciparum* immunogenic peptide fused to the C-terminus of MBP, was used. Cell proliferation was measured using the CellTiter 96® AQueous One Solution Cell Proliferation Assay (Promega, Madison, WI, USA) following the manufacturing protocol. Finally, plates were scanned at 490 nm with a Varioskan LUX (Thermo Fisher Scientific, Waltham, MA, USA) plate reader.

### Mouse tumor models and immunotherapy

Eight-week-old female C57BL/6 and BALB/c mice were purchased from Charles River Laboratories Italia (Lodi, Italy). Animal experiments were carried out in accordance with experimental protocols reviewed and approved (1060/2016-PR and 1153/2020-PR) by the Animal Ethical Committees of the University of Trento (Trento, Italy), Toscana Life Sciences Foundation (Siena, Italy) and by the Italian Ministry of Health. C57BL/6 and BALB/c mice were subcutaneously inoculated in the right flank with 1.5 × 10⁵ MC38 and with 2.5 × 10⁵ CT26, or 2.5 × 10⁵ WEHI-164 cells, respectively. When tumor size reached a volume between 70 and 100 mm³, mice received intratumoral injections of different OMV formulations (10 µg OMVs_Δ60_, 10 µg CCL3-OMVs_Δ60_, 10 µg FLT3L-OMVs_Δ60_, 5 µg IL-2-OMVs_Δ60_ + 5 µg TNFα-OMVs_Δ60_ Combo) in 50 µL PBS or PBS alone (50 µL). Treatments were administered three times at two-day intervals. Tumor growth was monitored for at least 60 days after the challenge using caliper measurements, and volumes were calculated as (A × B²)/2, where A is the longest and B the shortest diameter of the tumor. Statistical analysis of Kaplan-Meier curves was performed using Gehan–Breslow–Wilcoxon Test while unpaired, two-tailed Student’s *t*-test was used for the statistical analysis of tumor volume graphs (* p ≤ 0.05; ** p ≤ 0.01; *** p ≤ 0.001). Graphs were processed using GraphPad Prism 8 software.

### Flow cytometry analysis

Flow cytometry analysis of tumor cell populations was carried out as follows. CT26 tumors were collected and minced into pieces of 1–2 mm in diameter using a sterile scalpel and transferred into a 50 mL tubes. Then, the tumor tissue was enzymatically digested using the Tumor Dissociation kit (Miltenyi Biotech, Bergisch Gladbach, Germany) according to the manufacturer’s protocol, and the gentleMACS™ Dissociators (Miltenyi Biotech, Bergisch Gladbach, Germany) were used for the mechanical dissociation step. After tumor dissociation, the samples were filtered using a Cell Strainer 70 μm (Miltenyi Biotech, Bergisch Gladbach, Germany) followed by a second filtration step using a 30 μm filter (Miltenyi Biotech, Bergisch Gladbach, Germany) to remove larger particles and to collect a single-cell suspension. At the end of the dissociation protocol, 1 x 10^6^ cells (in 100 μL volume) were dispensed in each well of 96-well round plates and incubated with Fixable Viability Stain UV440 (BD Bioscience, San Jose, CA, USA) for 15 minutes at RT in the dark. Then, cells were washed twice in PBS and incubated with 25 µL of an anti-mouse CD16/CD32-Fc/Block (BD Bioscience, San Jose, CA, USA) for 20 minutes on ice in the dark. For each sample two independent staining protocols (A and B) were performed. A) 25 µL of the following mixture of fluorescent-labeled antibodies were added to the samples: CD3-APC (BioLegend, San Diego, CA, USA), CD4-BV510 (BioLegend, San Diego, CA, USA), CD8a-PECF594 (BD Bioscience, San Jose, CA, USA), CD25-PE-Vio770 (Miltenyi Biotech, Bergisch Gladbach, Germany), MHC II (I-Ek)-VioBright (Miltenyi Biotech, Bergisch Gladbach, Germany), CD11b-BV785 (BD Bioscience, San Jose, CA, USA) and Ly6C/Ly6G-BV421 (BD Bioscience, San Jose, CA, USA). B) 25 µL of the following mixture of fluorescent-labeled antibodies were added to the samples: CD3-APC (BioLegend, San Diego, CA, USA), CD4-BV510 (BioLegend, San Diego, CA, USA), CD8a-PECF594 (BD Bioscience, San Jose, CA, USA), TCRγδ-PE (Miltenyi Biotech, Bergisch Gladbach, Germany), TCRαβ-PEVio770 (Miltenyi Biotech, Bergisch Gladbach, Germany) and MHC II (I-Ek)-VioBright (Miltenyi Biotech, Bergisch Gladbach, Germany). In both protocols, cells were stained on ice in the dark for 30 minutes. After two washes with PBS, cells were fixed using Cytofix reagent (BD Bioscience, San Jose, CA, USA) for protocol B or with the Fixation/Permeabilization Solution of FoxP3 Staining Buffer Set (Miltenyi Biotech, Bergisch Gladbach, Germany) for protocol A. In this later case, after fixation cells were also incubated with an anti-FoxP3-PE (Miltenyi Biotech, Bergisch Gladbach, Germany) antibody diluted in permeabilization buffer and stained for 20 minutes at RT. Cells were finally washed twice in PBS (protocol B) or permeabilization buffer and then PBS (protocol A). Samples were analyzed using a Symphony A3 (BD Bioscience, San Jose, CA, USA) flow cytometry, and the raw data were elaborated using FlowJo software V10 6.1. For statistical analysis the unpaired, two-tailed Student’s *t*-test was used (* p ≤ 0.05; ** p ≤ 0.01; *** p ≤ 0.001) and graphs were processed using GraphPad Prism 8 software.

### Histology and Immunohistochemistry

Tumors were collected 24h after last injection, fixed for 20h at 4°C in buffered Formalin 10% (HT501128-4L, Sigma-Aldrich, St. Louis, MO, USA), processed with Histo-Line HistoPro200 Tissue Processor and embedded in paraffin with Histo-Line TEC 2900 Tissue Embedding Center. Five-micrometer thick sections were cut with the Leica HistoCore Biocut microtome and stained with hematoxylin and eosin (H&E) after deparaffinization and rehydration. Immunohistochemistry analyses were performed on deparaffinized sections following heat induced antigen retrieval performed with either EDTA pH9 buffer solution or antigen unmasking solution citrate-buffer (H3300, Vector Laboratories, Newark, CA, USA). Sections were kept 30 min at room temperature (RT) in blocking solution (PBS1X with 10% FBS), O/N at 4°C with primary antibody (anti-CD4, 1:1000 ab183685 Abcam, Cambridge, UK; anti-CD8, 1:500 ab217344 Abcam, Cambridge, UK; anti-MPO, 1:200 AF3667 R&D Systems, Minneapolis, MN, USA), and 2h at RT with biotinylated secondary antibody. Endogenous peroxidase was quenched in water diluted 3% H2O2 for 10 min at RT. Antibodies were visualized using an avidin-biotin complex (Vectastain Elite ABC kit, peroxidase PK-6100, Vector Laboratories, Newark, CA, USA) followed by diaminobenzidine (DAB peroxidase substrate kit SK-4100, Vector Laboratories, Newark, CA, USA) as a chromogen. Stained sections were analyzed with a Zeiss Axiolab 5 microscope equipped with an Axiocam 208 color camera and quantification was performed with QuPath software. Incubation with the secondary antibody and the avidin-biotin complex was used as control.

## List of supplementary materials

Figure S1 to S6

## Acknowledgments

We want to thank Simona Tavarini and Chiara Sammicheli for the highly professional support in flow cytometry analyses. We also express our gratitude to Silvia Valensin, Antonella De Rosa, Davide Bosco and Erika Bellini of the Animal Facility of Toscana Life Sciences Foundation (Siena, Italy) for the support in animal experiments. Finally, we thank Michela Roccuzzo (Advanced Imaging Core Facility, University of Trento, Italy) and Matteo Parri (Application Specialist High-End Microscopy, Nikon Europe B.V.) for the support in the confocal microscopy analysis.

## Funding

Advanced European Research Council grant VACCIBIOME (834634)

Proof-of-Concept ERC Grant INSITUOMVAC (101111780)

BiOMViS Srl (via Fiorentina, 1 Siena, Italy)

## Author Contributions

G.G. (Guido Grandi) and Al.G. (Alberto Grandi): development of the overall concept, design of the research and coordination of experimental activities; G.G., Al.G. and R.C.: writing of the manuscript; R.C., C.B., V.F., M.B., As.G. (Assunta Gagliardi), G.D.L. and Ga.G. (Gaia Gambini): plasmid cloning, bacteria-OMV preparations and analysis; G.D.L. and Al.G.: Dot Blot analysis; R.C., M.T. and M.B.: confocal microscopy analysis; R.C., E.C. and S.T.: cytokines *in vitro* activities; R.C., M.T., I.Z.: animal experiments and tumor challenge models; E.C, Al.G and R.C.: flow cytometry analysis on tumors; R.C.: immunofluorescence experiments and imaging analysis; R.B.: H&E staining and MPO immunohistochemistry on FFPE tumor sections; E.S.: 2DE gel preparation and 2DE Western Blot analysis; E.S., As.G. and L.B.: 2DE proteomics data analysis; R.C. and M.B.: figures preparation. A.L., A.B. and L.B. and ALL: critical review of published data and contribution to manuscript editing. All authors have read and agreed to the published version of the manuscript.

## Data and Materials availability

The data that support the findings of this study are available from the corresponding author upon reasonable request.

## Institutional Review Board Statement

The animal study protocol was approved by the Institutional Review Board (or Ethics Committee) of the University of Trento (Trento, Italy) and Toscana Life Sciences (Siena, Italy) and by the Italian Ministry of Health. (Protocols Code: 1060/2016-PR approved 9 November 2016 and 1153/2020-PR approved 23 November 2020).

