## Supplemental figures for "Cytokine-bearing Bacterial Outer Membrane Vesicles with Empowered Efficacy in Intratumoral Immunotherapy"

#### Supplementary figures

##### Figure S1

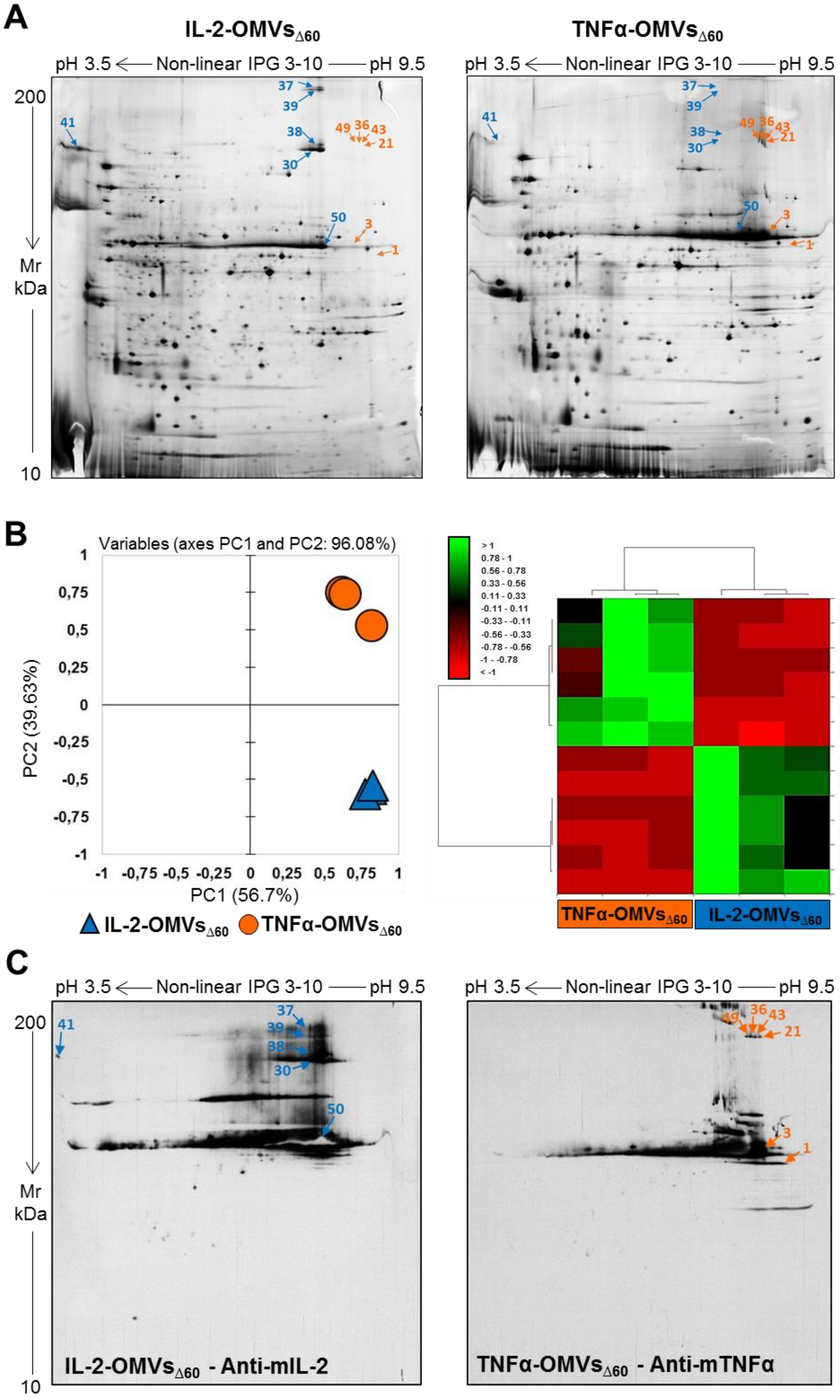

### Figure S2

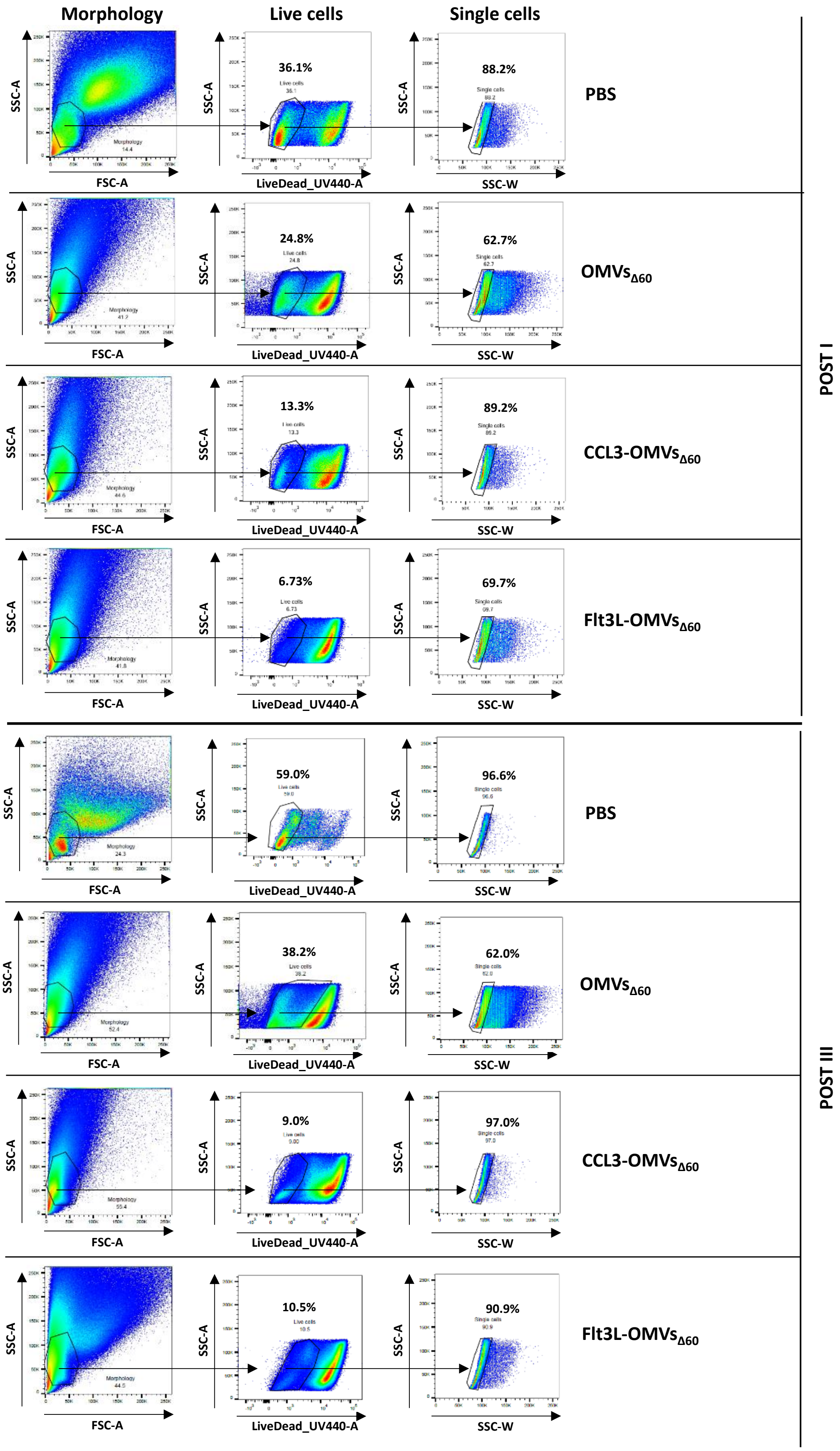
Morphology
Live cells
Single cells
SSC-A
FSC-A
LiveDead_UV440-A
SSC-W
36.1%
88.2%
PBS
24.8%
62.7%
OMVs$_{\Delta60}$
13.3%
89.2%
CCL3-OMVs$_{\Delta60}$
6.73%
69.7%
Flt3L-OMVs$_{\Delta60}$
POST I
59.0%
96.6%
PBS
38.2%
62.0%
OMVs$_{\Delta60}$
9.0%
97.0%
CCL3-OMVs$_{\Delta60}$
10.5%
90.9%
Flt3L-OMVs$_{\Delta60}$
POST III

### Figure S3

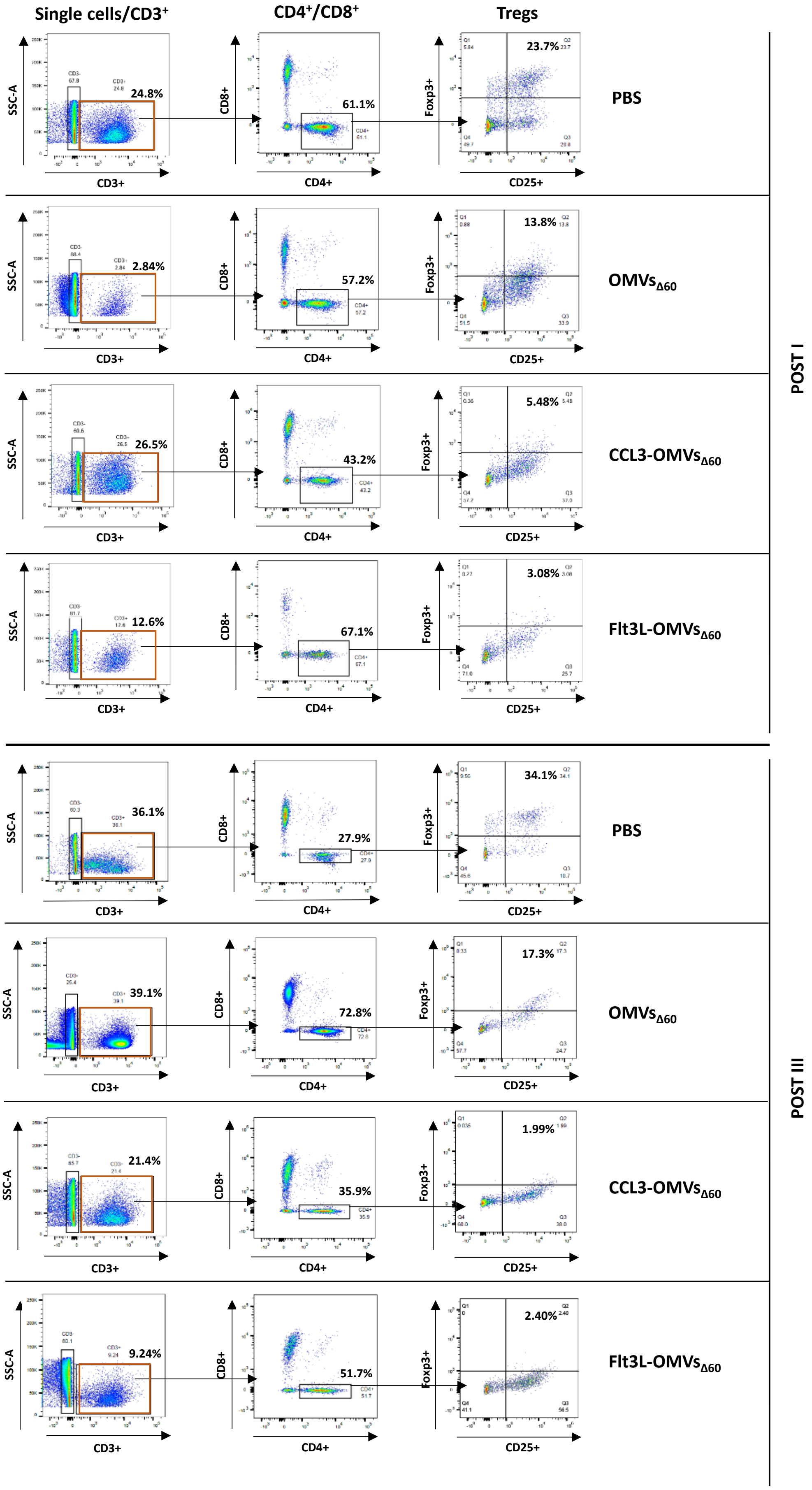

Single cells/CD3+
CD4+/CD8+
Tregs
PBS
OMVsΔ60
CCL3-OMVsΔ60
Flt3L-OMVsΔ60
POST I
POST III
SSC-A
CD3+
CD8+
CD4+
Foxp3+
CD25+
24.8%
61.1%
23.7%
2.84%
57.2%
13.8%
26.5%
43.2%
5.48%
12.6%
67.1%
3.08%
36.1%
27.9%
34.1%
39.1%
72.8%
17.3%
21.4%
35.9%
1.99%
9.24%
51.7%
2.40%

### Figure S4

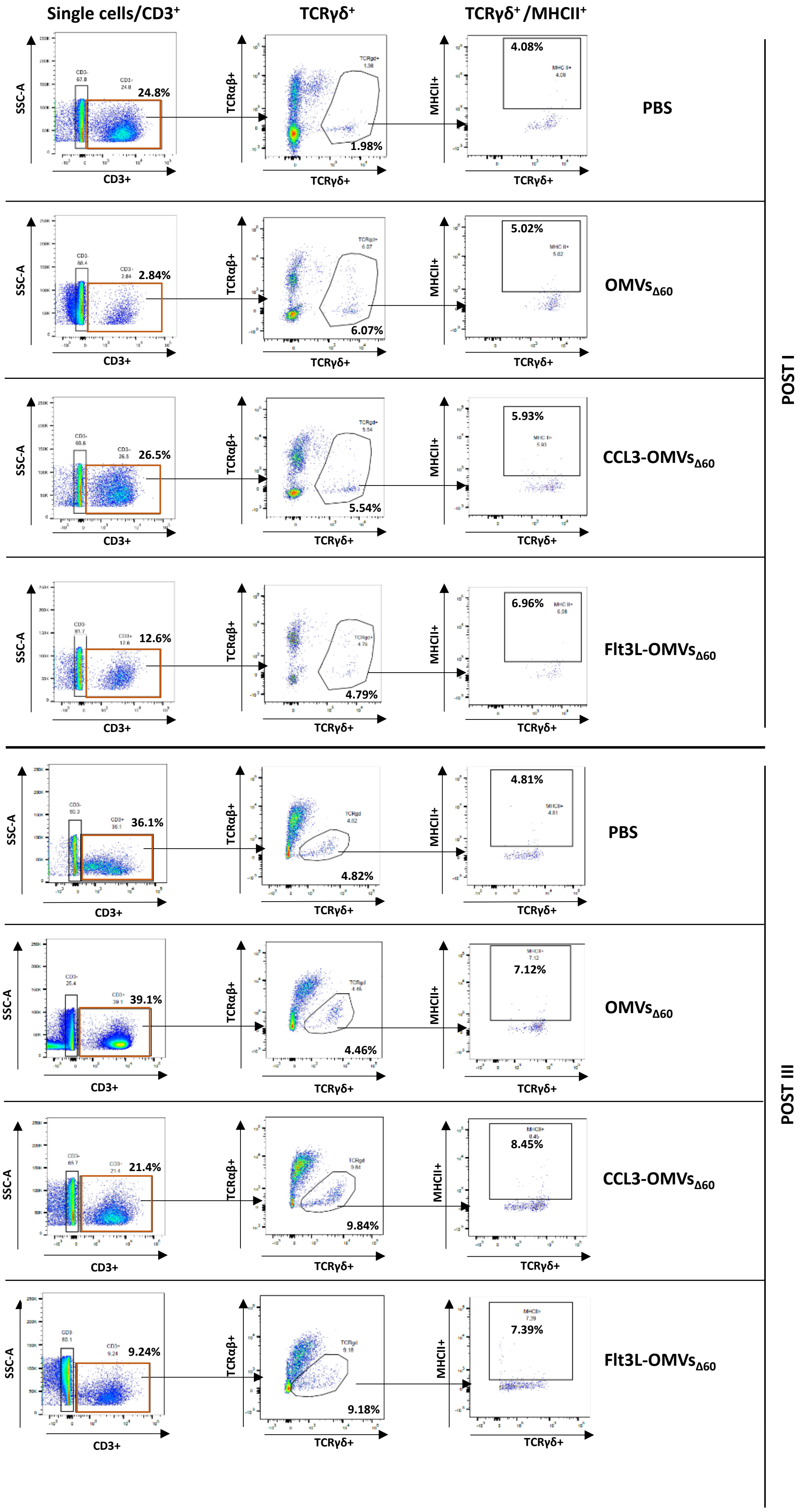

Single cells/CD3+
TCRγδ+
TCRγδ+ /MHCII+
PBS
OMVsΔ60
CCL3-OMVsΔ60
Flt3L-OMVsΔ60
POST I
POST III
SSC-A
CD3+
TCRαβ+
TCRγδ+
MHCII+
24.8%
1.98%
4.08%
2.84%
6.07%
5.02%
26.5%
5.54%
5.93%
12.6%
4.79%
6.96%
36.1%
4.82%
4.81%
39.1%
4.46%
7.12%
21.4%
9.84%
8.45%
9.24%
9.18%
7.39%

### Figure S5

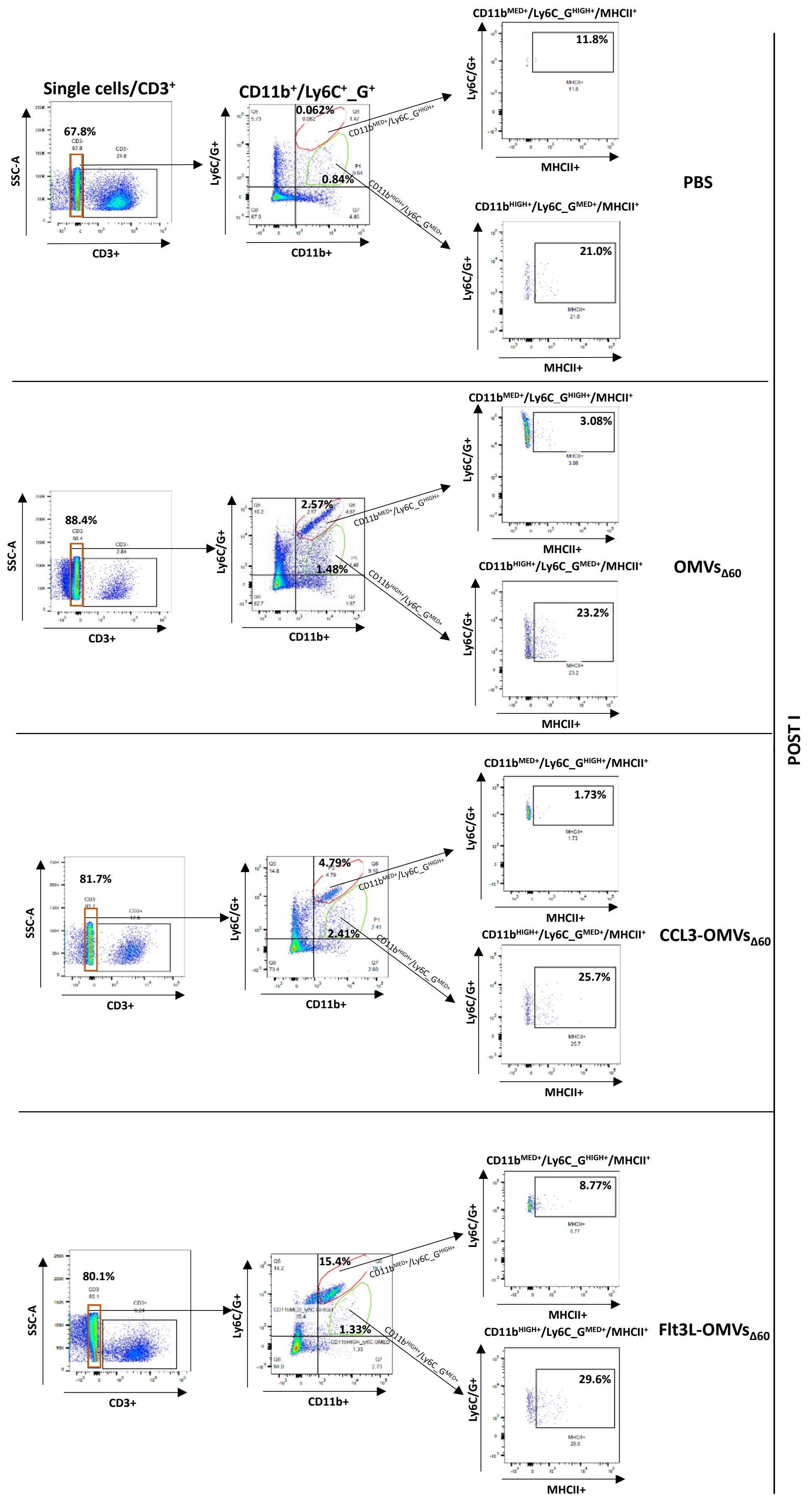

Single cells/CD3+
CD11b+/Ly6C+_G+
CD11bMED+/Ly6C_GHIGH+/MHCII+
CD11bHIGH+/Ly6C_GMED+/MHCII+
SSC-A
CD3+
Ly6C/G+
CD11b+
MHCII+
PBS
67.8%
0.062%
0.84%
11.8%
21.0%
OMVsΔ60
88.4%
2.57%
1.48%
3.08%
23.2%
CCL3-OMVsΔ60
81.7%
4.79%
2.41%
1.73%
25.7%
Flt3L-OMVsΔ60
80.1%
15.4%
1.33%
8.77%
29.6%
POST I

### Figure S6

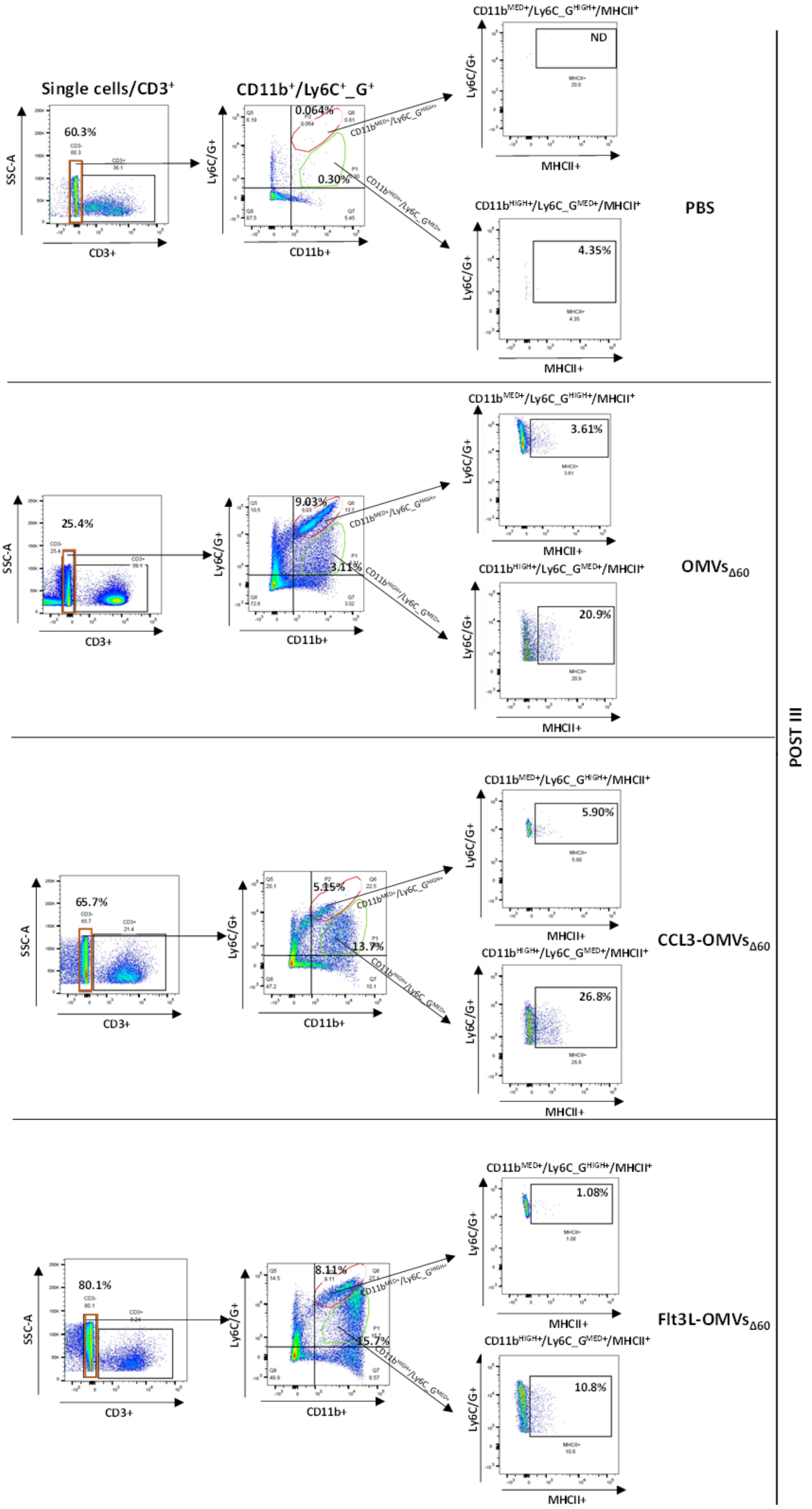
Single cells/CD3+
60.3%
SSC-A
CD3+
CD11b+/Ly6C+_G+
0.064%
0.30%
Ly6C/G+
CD11b+
CD11bMED+/Ly6C_GHIGH+/MHCII+
ND
MHCII+
CD11bHIGH+/Ly6C_GMED+/MHCII+
4.35%
PBS
25.4%
9.03%
3.11%
3.61%
20.9%
OMVsΔ60
65.7%
5.15%
13.7%
5.90%
26.8%
CCL3-OMVsΔ60
80.1%
8.11%
15.7%
1.08%
10.8%
Flt3L-OMVsΔ60
POST III

#### Supplementary figure legends

**Figure S1** – *Proteomic analysis* – Representative 2DE electropherograms of IL-2-OMVs$_{\Delta60}$ (left) and TNF$\alpha$-OMVs$_{\Delta60}$ (right). OMV proteins were properly denatured and separated by 2DE. 2DE gels were silver stained and the images acquired using the Image Scanner III laser densitometer equipped with LabScan 6.0 software (GE Healthcare). Image visualization and comparative analysis were carried out using dedicated software Melanie™ Classic 9 (SIB Swiss Institute of Bioinformatics, Geneva, Switzerland). Qualitative differential spots (ANOVA test; p-value ≤ 0.05; fold change ≥ 4.5) are marked with arrows and spot number, in blue for qualitative spots of IL-2-OMVs$_{\Delta60}$ and in orange for qualitative spots of TNF$\alpha$-OMVs$_{\Delta60}$ (**A**). Principal Component Analysis (PCA) of the qualitative differential spots obtained by 2DE image analysis between IL-2-OMVs$_{\Delta60}$ (blue triangles) and TNF$\alpha$-OMVs$_{\Delta60}$ (orange circles). PCA summarizes a 96.08% of variance (PC1: 56.63%; PC2: 39.45%) (**B, right**). Heatmap analysis of the qualitative differential spots obtained by 2DE image analysis between IL-2-OMVs$_{\Delta60}$ and TNF$\alpha$-OMVs$_{\Delta60}$. As shown in the legend, high abundant spots are reported in green, while low abundant spots are reported in red. Numbers on the right of the heat map refers to the spot numbers indicated in panel A (**B, left**). 2D-immunoblots of IL-2-OMVs$_{\Delta60}$ (left) and TNF$\alpha$-OMVs$_{\Delta60}$ (right) (**C**).

**Figure S2** – *Gating strategy applied to analyze tumor infiltrating immune cells* – BALB/c mice challenged with CT26 cells were intratumorally treated with either one or three doses (two days apart) of the following formulations: PBS (50 ml), OMVs$_{D60}$ (1 mg in 50 ml), CCL3-OMVs$_{D60}$ (1 mg in 50 ml), Flt3L-OMVs$_{D60}$ (1 mg in 50 ml) (tumor size at the time of treatment: approximately 100 mm$^3$). Twenty-four hours after treatments, tumors were collected and mechanically dissociated. Cells (1 x 10$^6$) were incubated with Fixable Viability Stain UV440 to define cell morphology and vitality. Only live, single cells were finally selected to characterize the populations of tumor-infiltrating immune cells. The Figure reports the flow cytometry analysis of tumors collected from one of the mice treated with either one dose (Post I) or three doses (Post III) of each formulation.

**Figure S3** – *Flow cytometry analysis of Regulatory T cells (Tregs) in tumors* – BALB/c mice challenged with CT26 cells were intratumorally treated with either one or three doses (two days apart) of the following formulations: PBS (50 ml), $OMVs_{D60}$ (1 mg in 50 ml), CCL3-$OMVs_{D60}$ (1 mg in 50 ml), Flt3L-$OMVs_{D60}$ (1 mg in 50 ml) (tumor size at the time of treatment: approximately 100 $mm^3$). Twenty-four hours after treatments, tumors were collected and mechanically dissociated. After applying the gating strategy shown in Figure S1, the viable single cells were analyzed after further surface staining with the following antibody mix: CD3-APC, CD4-BV510, CD8a- PECF594 and CD25-PE-Vio770. After Fixation/Permeabilization cells were also incubated with an anti-FoxP3-PE. The Figure reports the frequency of regulatory T cells (Tregs) present in tumors from one of the mice treated with either one dose (Post I) or three doses (Post III) of each formulation. $CD3^+$ cells (left panels) were first separated into $CD4^+$ and $CD8^+$ T cells (central panels). Then, the $CD4^+$ population was analyzed for the expression of CD25 and Foxp3 markers. The frequencies of the double-positive cells, corresponding to Tregs, are shown in the right panels of the figure.

**Figure S4** – *Flow cytometry analysis of γδ T cells in tumors* – BALB/c mice challenged with CT26 cells were intratumorally treated as described in Figure S1 legend. Twenty-four hours after treatments, tumors were collected and mechanically dissociated. The viable single cells (Figure S1) were analyzed after staining with the following antibody mix: CD3-APC, TCRgd-PE, TCRab-PEVio770 and MHC II(I-Ek)-VioBright. The Figure shows the frequencies of of γδ T cells present in tumors from one of the mice treated with either one dose (Post I) or three doses (Post III) of each formulation. CD3+ cells (left panels) were separated on the basis of the expression of either $\alpha\beta$ or $\gamma\delta$ T-cell receptor (central panels). The frequencies of $MHCII^+$, γδ T cells are shown in the right panels of the Figure.

**Figure S5** – *Flow cytometry analysis of myeloid cells in tumors after one vaccination* – BALB/c mice challenged with CT26 cells were intratumorally treated with one dose of the following formulations: PBS (50 ml), $OMVs_{D60}$ (1 mg in 50 ml), CCL3-$OMVs_{D60}$ (1 mg in 50 ml), Flt3L-$OMVs_{D60}$ (1 mg in 50 ml) (tumor size at the time of treatment: approximately 100 $mm^3$). Twenty-four hours after treatments, tumors were collected and mechanically dissociated. Viable single cells were analyzed after staining with the following antibody mix: CD3-APC, MHC II (I-Ek)-VioBright, CD11b-BV785 and Ly6C/Ly6G-BV421. The Figure shows the characterization of myeloid cells present in the tumors from one of the mice treated with each vaccine formulation. CD3 negative cells (left panels) were separated on the basis of the level of co-expression of Ly6C/Ly6G and CD11b protein markers. Two sub-populations were selected: $CD11b^{MED+}/Ly6C\_G^{HIGH+}$ (red gate) and the $CD11b^{HIGH+}/Ly6C\_G^{MED+}$ (green gate) (central panels). These two sub-populations were also analyzed for the expression of the MHC class II molecules (right panels).

**Figure S6 –** *Flow cytometry analysis of myeloid cells in tumors after three vaccination doses* – BALB/c mice challenged with CT26 cells were intratumorally treated with three doses (two days apart) of the following formulations: PBS (50 ml), $OMVs_{D60}$ (1 mg in 50 ml), CCL3-$OMVs_{D60}$ (1 mg in 50 ml), Flt3L-$OMVs_{D60}$ (1 mg in 50 ml) (tumor size at the time of treatment: approximately 100 $mm^3$). Twenty-four hours after treatments, tumors were collected and mechanically dissociated. Viable single cells were analyzed after staining with the following antibody mix: CD3-APC, MHC II (I-Ek)-VioBright, CD11b-BV785 and Ly6C/Ly6G-BV421. The Figure shows the characterization of myeloid cells present in the tumors from one of the mice treated with each vaccine formulation. CD3 negative cells (left panels) were separated on the basis of the level of co-expression of Ly6C/Ly6G and CD11b protein markers. Two sub-populations were selected: $CD11b^{MED+}/Ly6C\_G^{HIGH+}$ (red gate) and the $CD11b^{HIGH+}/Ly6C\_G^{MED+}$ (green gate) (central panels). These two sub-populations were also analyzed for the expression of the MHC class II molecules (right panels).
